# The distinct roles of MSH2 and MLH1 in basal-like breast cancer and immune modulation

**DOI:** 10.1101/2023.07.20.549745

**Authors:** Tanzia Islam Tithi, Jiao Mo, Nicholas Borcherding, Sung Jo, Heather R Kates, Edward Cho, Kailey E Cash, Masayoshi Honda, Lei Wang, Kawther K. Ahmed, Kalyanee Shirlekar, Li Chen, Katherine Gibson-Corley, Ronald Weigel, Maria Spies, Ryan Kolb, Weizhou Zhang

**Affiliations:** Department of Pathology, Immunology and Laboratory Medicine, College of Medicine, University of Florida, Gainesville, FL 32610, USA; Current: R & D, Thermo Fisher Scientific, Alachua, FL 32615, USA; Department of Pathology, the University of Iowa Carver College of Medicine, Iowa City, IA, 52242; Current: Department of Pathology and Immunology, Washington University School of Medicine, St Louis, MO 63110, USA. Electronic address; Current: R & D, Carbon Biosciences, Waltham, MA 02451; Department of Surgery, the University of Iowa Carver College of Medicine, Iowa City, IA, 52242; Department of Biochemistry and Molecular Biology, the University of Iowa Carver College of Medicine, Iowa City, IA, 52242; Current: Department of Pharmaceutics, the University of Baghdad College of Pharmacy, Bab-almoadham, PO Box 14026, Baghdad, Iraq; University of Florida Health Cancer Center (UFHCC), the University of Florida, Gainesville, FL 32610, USA; Department of Biostatistics, College of Medicine, University of Florida, Gainesville, FL 32610, USA; Department of Pathology, Microbiology and Immunology, Vanderbilt University Medical Center, Nashville, TN 37232

## Abstract

The mismatch repair (MMR) pathway is known as a tumor suppressive pathway and genes involved in MMR are commonly mutated in hereditary colorectal or other cancer types. However, the function of MMR genes/proteins in breast cancer progression and metastasis are largely undefined. We found that MSH2, but not MLH1, is highly enriched in basal-like breast cancer (BLBC) and that its protein expression is inversely correlated with overall survival time (OS). *MSH2* expression is frequently elevated due to genomic amplification or gain-of-expression in BLBC, which results in increased MSH2 protein to pair with MSH6 (collectively referred to as MutSα). Genetic deletion of *MSH2* or *MLH1* results in a contrasting phenotype in metastasis, with *MSH2*-deletion leading to reduced metastasis and *MLH1*-deletion to enhanced liver or lung metastasis. Mechanistically, MSH2 – but not MLH1 – binds to the promoter region of interferon α receptor 1 (*IFNAR1*) and suppresses its expression in BLBC. Deletion of MSH2 initiates a chain of immune reactions via the upregulation of IFNAR1 expression and the activation of type 1 interferon signaling, which explains a highly immune active tumor microenvironment in tumors with MSH2-deficiency. Our study supports the contrasting functions of MSH2 and MLH1 in BLBC progression and metastasis due to the differential regulation of IFNAR1 expression, which challenges the paradigm of the MMR pathway as a universal tumor suppressive mechanism.

## Introduction

Basal-like breast cancer (BLBC) is an aggressive molecular subtype of breast cancer, accounting for 12.3-36.7% of invasive ductal carcinomas in different patient cohorts(Toft and Cryns, 2011). The absence of approved targeted therapy and the variable responses to chemotherapy in BLBC patients represent vital and unmet clinical need. BLBC is also defined by high levels of genomic instability, with p53 pathway alterations seen in 85-95% of tumors (Toft and Cryns, 2011). This instability in BLBC is compounded by the characteristic loss in chromosome 5q, consisting of DNA repair genes, *RAD17*, *RAD50*, *MSH3*, and X*RCC4 (Rakha et al., 2008).* In addition,15-20% of BLBC possess mutations in *BRCA1* or *BRCA2,* which function in double-stranded DNA repair. Half of the 15-20% of BLBC tumors with *BRCA1* or *BRCA2* are a result of germline mutations and are associated with a 70% lifetime risk of breast or ovarian cancer to individuals (Rakha et al., 2008). Recent success of clinical trials using inhibitors for Poly-ADP-ribose polymerase 1 (PARP1), a single-strand DNA repair sensor, in germline *BRCA*-mutated ovarian and breast cancers have led to Food and Drug Administration approval(Zimmer et al., 2018). PARP1 plays multiple role in genome maintenance, signaling and repair of DNA damage and stabilization of stalled DNA replication forks (Lord and Ashworth, 2017; Ray Chaudhuri and Nussenzweig, 2017). Its pharmacological inhibition leads to cell death in cells carrying biallelic BRCA mutations (Ashworth, 2008; Hengel et al., 2017; Zimmer et al., 2018). Despite the positive successes of PARP inhibitors in trials, the translation of PARP therapy to the majority of BLBC and ovarian cancers with intact *BRCA1/2* have been unsuccessful. Even in patients with BRCA-mutated breast cancers, 5-year survival with Olaparib is about 15%, and that the rest develop resistance (Incorvaia et al., 2017; Ledermann et al., 2016; Lupo and Trusolino, 2014; Murata et al., 2016).

The success of PARP inhibitors for a subset of BLBC/TNBC patients has motivated the field to investigate the potential of other DNA repair-based targeted therapy. Mismatch repair (MMR) genes have traditionally been thought of as tumor suppressors, with germline mutations identified in the autosomal dominant Lynch Syndrome (Bronner et al., 1994; Fishel, 2015; Fishel et al., 1993; Leach et al., 1993). Closely associated with hereditary non-polyposis colorectal cancer, Lynch syndrome also increases risk in the development of cancers along the gastrointestinal tract, endometrial cancer, and ovarian cancer (Modrich, 1994). Defective MMR (dMMR) genes, including MLH1, MSH2, MSH6 and PMS2, lead to the increase in mutations and are commonly associated with microsatellite instability (MSI) (Leach et al., 1993; Parsons et al., 1993) and mutator phenotypes (Drummond et al., 1995; Modrich, 1994). This increased mutational rate is thought to underlie the 25-80% therapeutic response rate of anti-PD-1 immune checkpoint inhibitor (ICI) in patients with Lynch-Syndrome-associated tumors(Geoerger et al., 2020; Le et al., 2020; Le et al., 2015; Marabelle et al., 2020). The result of this trial led to the unprecedented approval of anti-PD-1 therapies for metastatic dMMR or MSI cancers, irrespective of the site of origin. Interestingly, patients with MSI or dMMR exhibit very different responses to ICIs, depending on the extent of insertion-deletion mutations (Mandal et al., 2019) or the status of cGAS-STING-dependent DNA sensing and interferon β (IFN-β) activation (Guan et al., 2021), leading to the recruitment of CD8^+^ T cells by CCL5 and CXCL10 production (Mowat et al., 2021). Though most of these studies are based on genetic silencing or knockout of key MMR genes *MLH1* or *MSH2*, those two act epistatically and are commonly considered equal and have similar impacts on cancer progression or immune regulations via the MMR pathway.

Despite acting epistatically in the mismatch recognition and performing tumor suppressor functions, we found that MSH2 and MLH1 have contrasting roles in the metastasis of BLBC. MSH2 protein is associated with decreased immune infiltration and increased metastasis; whereas MLH1 suppresses metastasis. MSH2-deletion promotes immune cell infiltrations via inducing a systematic increase in chemokines by potentiating IFNAR1-mediated signaling transduction, following with downstream induction of immune related pathways. This study identifies the opposite roles of different MMR proteins in cancer progression and metastasis and establishes the rationale to target MSH2 or MutSα for BLBC therapy.

## Results

### MSH2 and MSH6 are elevated in BLBC and predict poor overall survival

We previously developed analysis workflow to identify survival outcomes using reverse-phase protein array (RPPA) (Li et al., 2013) quantification available in the Cancer Genome Atlas (TCGA) (Borcherding et al., 2018). We applied the system to examine prognostic markers for overall survival in BLBC (Fig. 1A), with a focus on DNA repair proteins available on the RPPA (highlighted in red). Of the 6 DNA repair proteins that had significant (*P* < 0.05) prognostic value, elevated expression of MSH2, RAD50, KU80, CHK2, and MSH6 predicted poor overall survival with Cox proportional hazard ratios greater than 1 (Fig. 1B). The division of samples in high versus low was based on maximal standardized two-sample linear rank statistic to find an optimal cutpoint, providing a range proportional comparison (Fig. 1C). Notably, MSH2 evenly split at 50% of samples (Fig. 1C) and predicted the worse survival for any DNA repair protein (Fig. 1D, upper panel, hazard ratio=8.2). After removing unfollowed patients from the BLBC cohorts, MSH6, which forms a heterodimer with MSH2, had a same optimal cutpoint at 50%, and predicted poor overall survival (Fig. 1D, lower panel, hazard ratio=7.6). Since the TCPA dataset includes limited proteins, we also expanded the analysis using the TCGA RNA sequencing data where we found that *MLH1* and *PMS1* (collectively referred to as MutLα) expression predicts better prognosis with HR of 0.65 and 0.42 respectively (Supplementary Table 1). Other poor prognosis predictors in the DNA repair pathways include *MSH3* in the MMR pathways, as well as *MPG* and *NTHL1* in the base excision repair (BER) pathway (Supplementary Table 1). We also performed Cox regressions in the other molecular subtypes and found that there is no significant predictive value for MSH2 and MSH6 in HER2+ or Luminal A breast cancers (Supplementary Fig. 1). Interestingly, in luminal B breast cancer, MSH2 protein level predicted improved overall survival (Supplementary Fig. 1). Across breast cancer samples, MSH2 and MSH6 protein levels were directly correlated with one another (Fig. 1E), supporting the established literature on the need of heterodimer formation for MSH6 stabilization (Arlow et al., 2021). Accompanying the survival predictions, we also found MSH2 and MSH6 (Fig.1F) are significantly elevated in BLBC compared to the other breast cancer molecular subtypes. MSH3 pairs with MSH2 to form MutSβ, but MSH3 is excluded in these bioinformatics analysis because gene locus of MSH3 is frequently deleted in upto 70% TNBC/BLBC (Johannsdottir et al., 2006; Natrajan et al., 2009; Turner et al., 2010).

**Figure 1.**
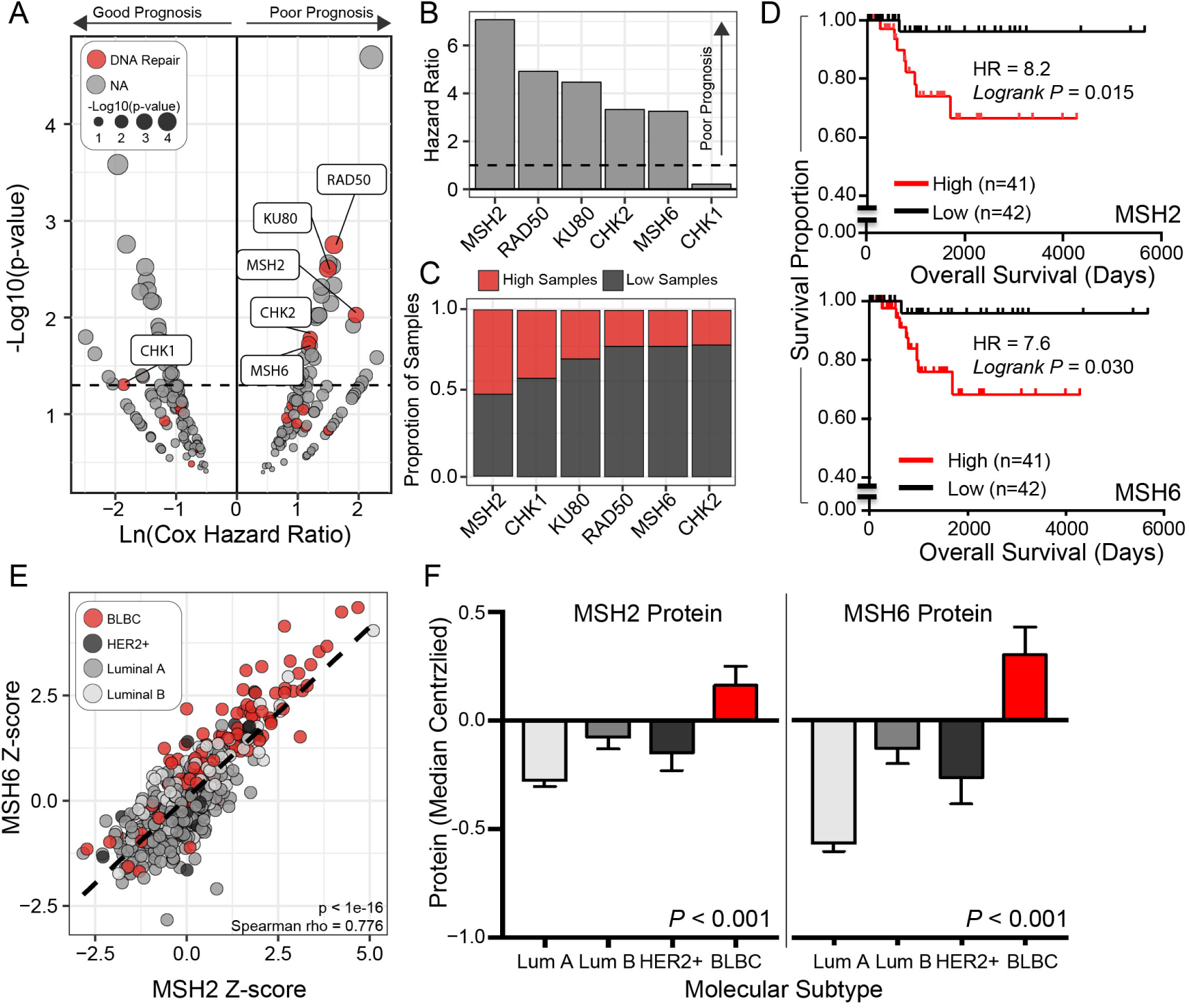
DNA mismatch repair MSH2 protein is elevated in BLBC and predicts poor survival. **A.** Cox regression hazard ratio and p-values across all 224 proteins on the RPPA for 119 BLBC samples. DNA repair proteins are highlighted in red. **B.** Hazard ratio (HR) summary for the 6 RPPA proteins with a *P* value less than 0.05. HR above one (dotted line) indicates poor survival for the higher protein level. **C.** Proportional cut points for the six significant RPPA proteins based on log-rank optimal *P* value. **D.** Survival curve for MSH2 (higher panel) or MSH6 (lower panel) in 83 BLBC samples comparing protein-high (red, n=41) to protein-low (black, n=42) specimens in the TCGA dataset with unfollowed cases removed from the analysis. Logrank test was used and *p* values indicated. **E.** MSH2-MSH6 protein correlations for breast cancer samples with molecular subtype designations (n=670, *P* < 1e-16). **F.** MSH2 or MSH6 RPPA protein quantification across molecular subtypes, BLBC (n=119), HER2+ (N=61), Luminal A (n=328), and Luminal B (n=162). RPPA protein data from the TCPA was analyzed, median-centralized and compared. One-way ANOVA was used with *p* values indicated.

### The contrasting roles of MSH2 and MLH1 in breast cancer metastasis

The differential impacts of MSH2 and MLH1 on BLBC prognosis indicate the potential different functions of them in BLBC progression. We further expanded the analysis among the TCGA pan-cancer datasets and found that *MSH2* is frequently amplified with the gain of mRNA expression in breast cancer (BRCA), lung cancers (LUAD and LUCA), ovarian cancer (OV) *etc* (Supplementary Fig. 2A). MLH1, in contrast, exhibits frequent mutations and deletions but with few amplification or gain of expression (Supplementary Fig. 2B), further supporting the potentially different functions of MSH2 and MLH1 among different cancer types.

To determine the roles of MSH2 and MLH1 in BLBC, we used protein-gRNA (RNP) transfection-based CRISPR/Cas9 system to knock out their expression in several BLBC cells, including murine 4T1 (Supplementary Fig. 3A-B) and Py8119 (Supplementary Fig. 3C-D) breast cancer cells, murine MC38 colon cancer cells (Supplementary Fig.4A-B), and human MDA-MD-231 (Supplementary Fig. 4C-D) basal B breast cancer cells. All single clones were validated for the deletion of target genes and several clones were combined (at least 5 knockout clones, referred to as either control (Con) or knockout (KO)) for each cell type (Supplementary Fig. 3-4) for tumor and metastasis studies (Fig. 2, Supplementary Fig. 5). MSH2 KO did not significantly influence primary tumor growth from Py8119 (Fig. 2A), 4T1 (Fig. 2C), MC38 (Supplementary Fig. 5A) or MDA-MB-231 (Supplementary Fig. 5B), but led to a reduction of lung metastasis in Py8119 (Fig. 2B, *P* = 0.009) or 4T1 (Fig. 2D, 3/7 mice with 4-5 metastatic nodules in control group while no mice in the MSH2 KO group) orthotopic models, or lung and liver experimental metastasis via *i.v.* injection of Py8119 cells in immune competent C57BL/6J (Py8119) or Balb/C mice (4T1 model) (Fig. 2E-F). MLH1 KO, however, led to an increase in primary tumor growth and metastasis in the Py8119 and 4T1 models (Fig. 2B, 2D), but not in the MC38 colon cancer model (Supplementary Fig. 5A). The *i.v.* experimental metastasis model further validates the significant increase in metastasis due to *MLH1* KO, indicating that tumor size increase may not be the primary reason of increased metastasis due to MLH1 deficiency. It is worth noting that the MSH2-mediated metastasis relies on the intact immune system as *i.v.* injection of Py8119 cells into the NOD/Scid/IL-2Rγ^-/-^ (NSG) mice – lacking functional T, B and nature killer (NK) cells – had equal amount of metastasis between control and MSH2 KO groups (Fig. 2G). When injecting the human MDA-MB-231 cells into NSG mice, MLH1 KO also led to increase in tumor growth and tendency of shorter overall survival of the mice (Supplementary Fig. 5B-C), suggesting that the MLH1-mediated tumor/metastasis-suppressing phenotypes are likely via cancer-cell intrinsic pathways and are independent of T, B or NK cells.

**Figure 2.**
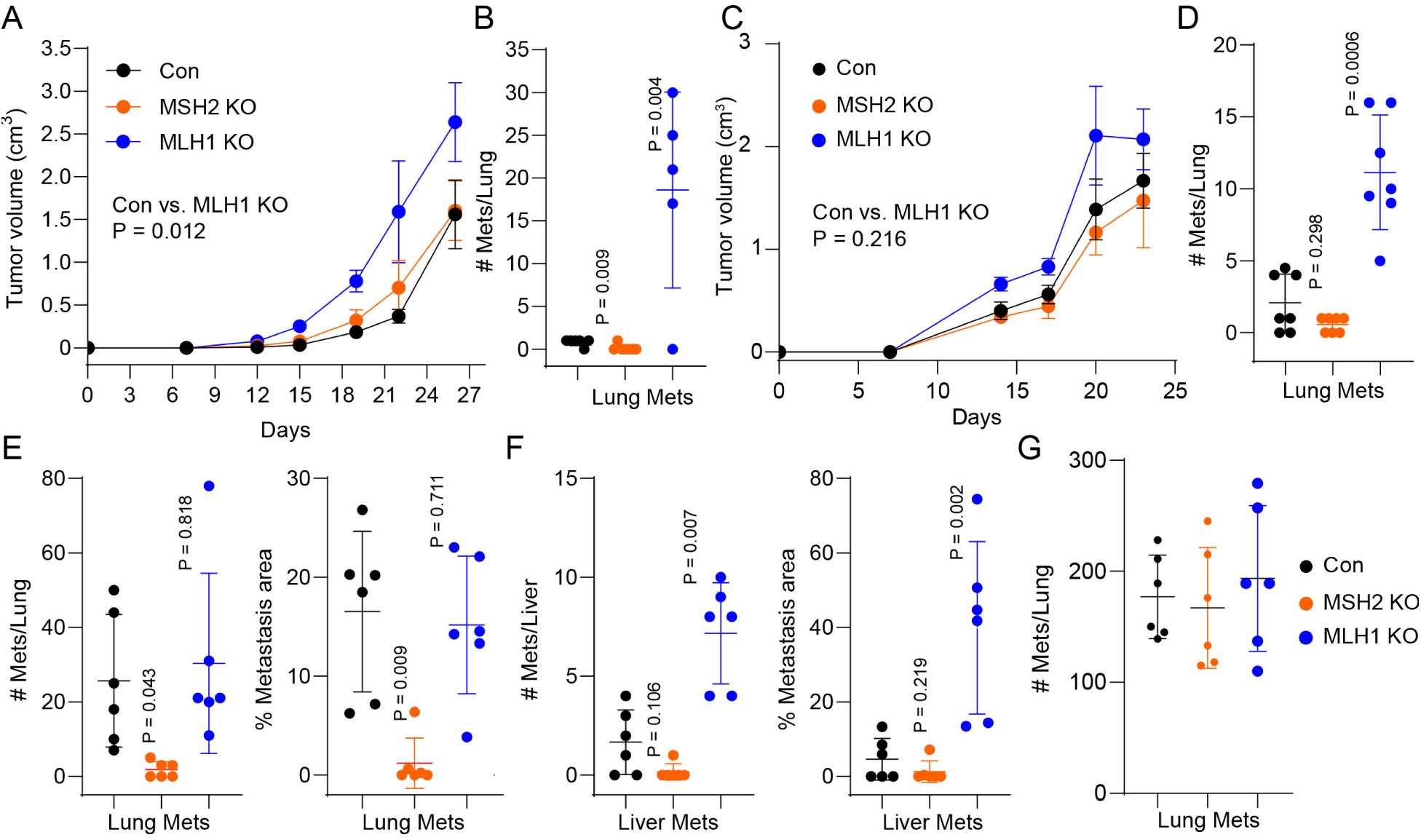
Opposing functions of MSH2 and MLH1 in breast cancer metastasis. **A-B.** Primary tumor growth curve (A) or lung metastasis (B) of control (Con), MSH2-KO or MLH1-KO cells of Py8119 orthotopically injected into #4 fatpad of in 8-week old female C57BL/6J (n = 7 per group). **C-D.** Primary tumor growth curve (C) or lung metastasis (D) of control (Con), MSH2-KO or MLH1-KO cells of 4T1 orthotopically injected into #4 fatpad of 8-week old female Balb/C mice (n = 7 per group). **E-F.** Lung (E) or liver (F) metastasis of Con, MSH2-KO or MLH1-KO cells of Py8119 *i.v.* injected into 8-week old female C57BL/6J (n = 7 per group). **G.** Lung metastasis of Con, MSH2-KO or MLH1-KO cells of Py8119 *i.v.* injected into 8-week old female NSG mice (n = 7 per group).

### The differential roles of MSH2 and MLH1 in tumor-infiltrating (TI) immune cells in BLBC

As deletion of cancer cell-intrinsic MSH2 or MLH1 has opposite phenotypes in metastasis and that MSH2-dependent tumor/metastasis promotion relies on an intact immune system, we determined the immune cell compositions as we established before (Kolb et al., 2021), including tumors or spleens (Supplementary Fig. 6A-B, Fig. 3) of the Py8119-tumor bearing mice (Fig.2A). MSH2 KO in Py8119 cells did not significantly change the peripheral immune profiles (spleen) in general (Fig. 3A, 3C, 3E, 3G, 3I, 3K, 3M), except for the slight decrease in dendritic cells (DC, CD11c^+^MHCII^+^) and active DCs (CD80^+^) (Fig. 3I), or monocytes (CD11b^+^Ly6C^+^Ly6G^-^) (Fig. 3M). In contrast, MSH2 KO led to significantly increase in majority of TI immune cells, including total leukocyte or lymphocytes (Fig. 3B), NK (CD3^-^NK1.1^+^, active NK (Granzyme B^+^), NKT (CD3^+^NK1.1^+^) (Fig. 3D), conventional CD4^+^ T cells (Tconv, CD4^+^Foxp3^-^), regulatory T cells (Treg, CD4^+^CD25^+^Foxp3^+^) (Fig. 3F), CD8^+^ and active CD8^+^ (Granzyme B^+^/Perforin^+^) T cells (Fig. 3H), neutrophils (Fig. 3L) or monocytes (Fig. 3N), without significant changes in tumor associated macrophages (TAM, Fig. 3N).

**Figure 3.**
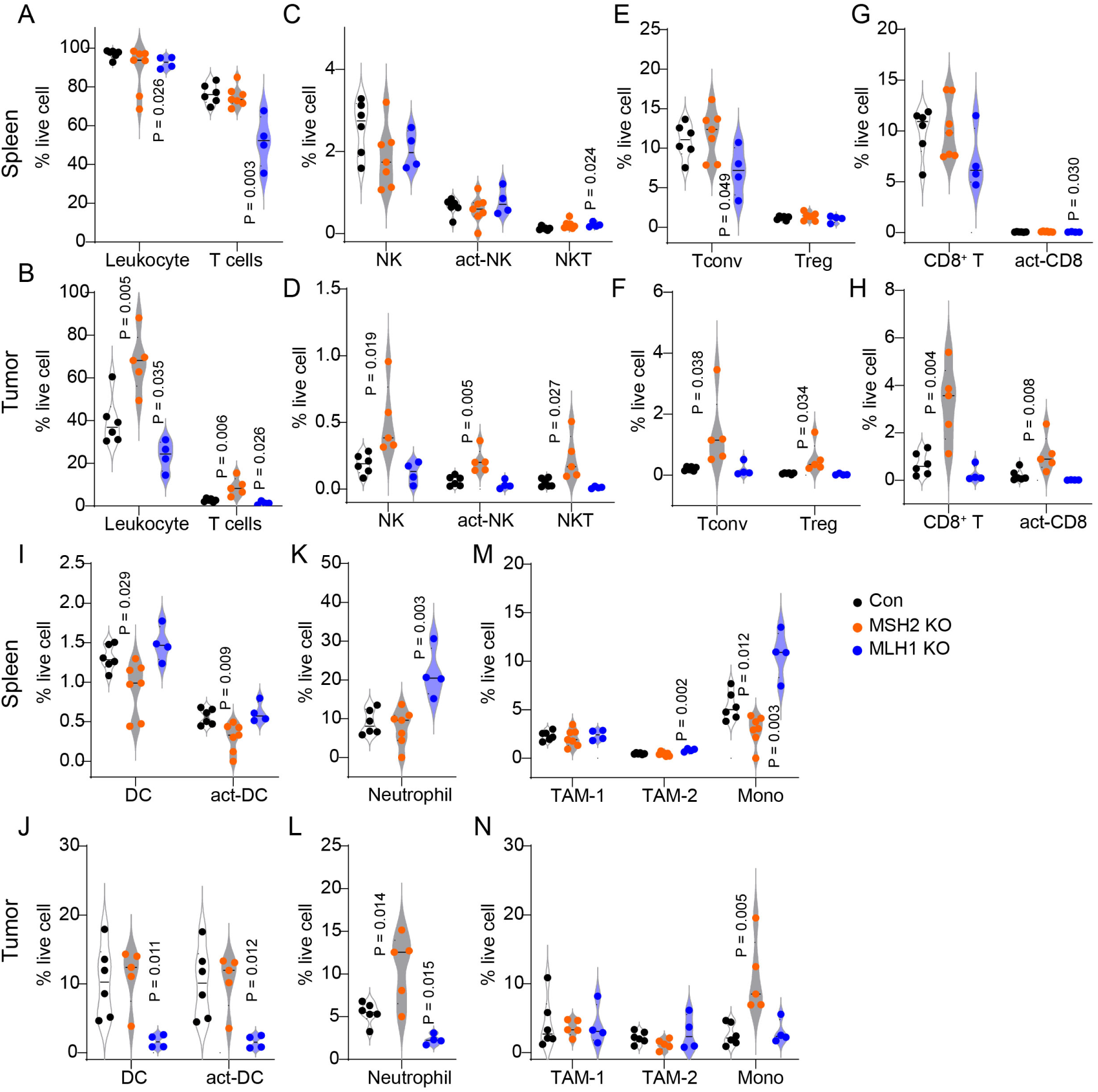
MSH2 KO leads to an overall increase in tumor-infiltrating leukocytes. Tumors and spleens from Fig. 2A were analyzed for immune profiles including (A-B) CD45+ leukocytes; (C-D) CD3+ T cells; (C) NK1.1+ NK cells, NK1.1+GZMB+ active NK cells, or CD3+NK1.1+ NKT cells; (E-F) CD3+CD4+Foxp3-conventional T cells (Tconv) or CD3+CD4+Foxp3+ Treg cells; (G-H) CD3+CD8+ T cells or CD3+CD8+GZMB+ active CD8 T cells; (I-J) CD11c+MCHII+ dendritic cells or CD86+ DCs (act-DC); (K-L) CD11b+Ly6G+ neutrophils; and (M-N) two distinct tumor associated macrophages (CD11c+F4/80+ and CD11c-F4/80+) or monocytes (CD11b+Ly6C+). N = 4-5 per group.

We also determined the immune cell infiltrations including tumors or spleens of the 4T1-tumor bearing mice (Supplementary Fig. 6C-D), including mixed control clones (con-mix), mixed MSH2 KO clones (KO-mix), as well as individual MSH2 KO clones. In agreement data shown above (Fig. 2A,2C), MSH2 KO did not significantly influence primary tumor growth (Supplementary Fig. 6C), but induced significant CD45^+^ immune cell infiltrations in the primary tumors of but not in spleen (Supplementary Fig. 6D).

MLH1 KO, interestingly, has an impact more on peripheral immune profiles than on TI immune profiles (Fig. 3A, Fig. 3K), with a marked decrease in peripheral lymphocytes (Fig. 3A) and a dramatic increase in neutrophils (Fig. 3K, an increase from 9% in the control group to 21% in the KO group, *P*=0.003) and monocytes (Fig. 3N, *P*= 0.003). Other altered cells in the periphery included a slight increase in NKT cells (Fig. 3C) and minor decrease in active CD8^+^ T cells (Fig. 3G). In the TME, MLH1 KO led to significant decreases in total leukocytes and lymphocytes (Fig. 3B), DCs, active DCs (Fig. 3J), and neutrophils (Fig. 3L), without significant influences on NK, T, or TAM subpopulations (Fig. 3D, 3F, 3H). These immunological changes induced by deletion of cancer cell intrinsic MSH2 or MLH1 agree with the tumor/metastasis phenotypes observed in BLBC tumors (Fig. 2).

### Differential effects of MSH2- or MLH1-KO on the chemokine/cytokines and immune activation in BLBC

To understand the differential roles of MSH2 and MLH1 in immune modulation, we performed RNA sequencing of total RNAs from tumors in Fig. 2A and Fig. 2C. Gene Set Enrichment Analysis (GSEA) identified that the chemokine pathways are the only consistent pathways that are enriched in MSH2 KO Py8119 (Fig. 4A) and 4T1 (Fig. 4B) tumors, relative to both control and MLH1 KO tumors (Fig. 4C), in agreement with the general increase in TI immune cell populations (Fig. 3). Other pathways involved in immune modulations include a panel of MHC molecules in the Py8119 MSH2 KO tumors (Fig. 4D), in agreement with immune active phenotype and T cell recognition. MSH2 KO in BT549 cells led to a similar MHC upregulation, indicating an activation of antigen presentation pathways via cancer-intrinsic mechanisms (Supplementary Fig. 7). We collected conditioned medium from WT 4T1 cells, or MSH2 KO or MLH1 KO 4T1 cells, and found that several cytokine/chemokine proteins were also selectively elevated in the MSH2 KO cells relative to WT and MLH1 KO cells (Fig. 4E, Supplementary Fig. 8), suggesting the cancer-cell intrinsic changes in chemokine/cytokine production mediated by MSH2.

**Figure 4.**
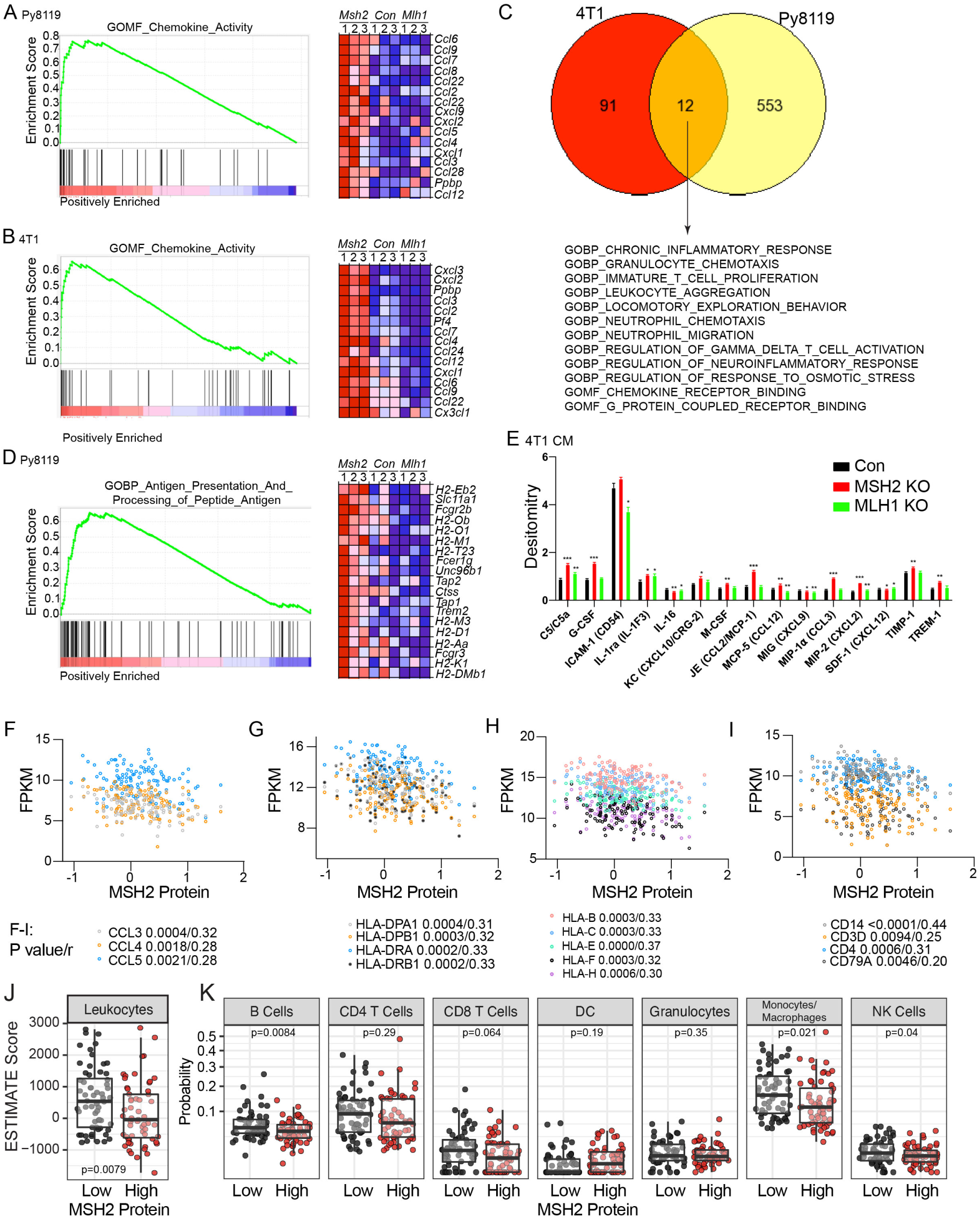
MSH2 KO induces immune active tumor microenvironment via chemokine induction. **A-D**. Tumors formed from parental (Con), MSH2-KO (*Msh2*) or MLH1-KO (Mlh1) cells of Py8119 (Fig. 2A) or 4T1 (Fig. 2C) were subject to RNA sequencing, following with GSEA analysis using the C5: ontology gene sets. MSH2-KO tumors enrich in pathways related to chemokine activity in (A) Py8119 or (B) 4T1 BLBC model related to Con and MLH1-KO tumors; (C) Venn diagram showing the 12 common ontology genesets in the MSH2-KO tumors of both Py8119 and 4T1 models; and (D) MSH2-KO tumors enrich in pathways related to antigen presentation. **E.** MSH2-KO increases immune modulatory chemokine/cytokine proteins in conditional medium using Proteome Profiler Mouse Cytokine Array Kit, Panel A (Biotechne, R&D). **F-I.** Correlation of MSH2 protein with immune modulatory molecules within the TCGA Breast Cancer BLBC dataset, including (F) chemokines; (G) MHC II genes; (H) MHC I genes; (I) immune cell lineage markers. **J-K.** Immune infiltration score based on the ESTIMATE algorithm by MSH2 mean protein value, comparing MSH2-high (n=60) to MSH2-low (n=59) BLBC specimens from the TCGA breast cancer dataset. **(J)** total ESTIMATE score for all leukocytes (P value is determined using Welch’s t-test); (**K)** Condensed CIBERSORT estimates of immune cell lineages based on mRNA values available for the BLBC samples.

Datamining the TCGA dataset, we found that MSH2 protein is significantly and inversely correlated with chemokines such as CCL3, CCL4, CCL5 (Fig. 4F, Supplementary Table 2), MHC molecules (Fig. 4G-H, Supplementary Table 2) and markers for monocytes, T cells or B cells (Fig. 4I, Supplementary Table 2) among the human BLBC patient specimens. In contrast, MLH1 expression was positively correlated with immune-related genes or immune cell markers in human TCGA BLBC specimens (Supplementary Table 3), further supporting the contrasting effects of MSH2 and MLH1 in immune regulations within BLBC cancer type. In addition, we separated human BLBC specimens by MSH2 protein level (high versus low) and estimated the immune profiles using ESTIMATE and CIBERSORT algorithms. Consistent with the correlation studies (Fig. 4I), we found increased estimated immune cell infiltration (Fig. 4J), with lineage specific increases in B cells (*P*=0.0084), monocytes/macrophages (*P*=0.021), NK cells (*P*=0.04), and CD8 T cells approaching significance (*P*=0.062) (Fig. 4K). GSEA analysis further revealed that MSH2-low BLBC specimens have significantly enriched signaling pathways involved in immune regulations (Supplementary Table 4); whereas MSH2-high BLBC specimens have elevated pathways including RNA processing, damaged DNA binding, microtubule organization, DNA strand elongation, chromosomal segregation *etc*. that are important for cancer cell proliferation and progression (Supplementary Table 5). In contrast, MLH1-low BLBC specimens have elevated pathways involved in ribosomal functions, translation, mitochondrial functions *etc* (Supplementary Table 6, Supplementary Fig.9A-D), indicating more active and aggressive cancers, which are in line with the results shown in Fig. 2. MLH1-high BLBC specimens did not exhibit any distinct pathways related to cancer progression (Supplementary Table 7).

We performed differential gene expression compared MSH2-high (n=60, red) to MSH2-low (n=59, black) using estimated counts (Supplementary Fig. 10A). As expected, we found MSH2-high associated with increased RNA levels of *MSH2* and *MSH6*, in addition to increased *PMS1*, which participates in the repair downstream of the MutSα heterodimer (Supplementary Fig. 10B). MSH2-low BLBC samples had increased levels of immune checkpoints, *IDO1*, *CD274*, and *HAVCR2* (Supplementary Fig. 10C), indicating an immune active but exhausted tumor microenvironment. This increased levels of immune markers in MSH2-low samples was seen across chemokines/cytokines (Supplementary Fig. 10D) and chemokine/cytokine receptors (Supplementary Fig. 10E). We also performed Ingenuity Pathway Analysis (IPA) for upstream regulators of the transcriptional differences observed (Supplementary Fig. 10F). Quadrant I refers to upstream regulators that have increased RNA expression and activation in MSH2-high BLBC samples, while quadrant III refers to regulators with increased expression and activation in MSH2-low samples. In MSH2-low BLBC samples, we observed a general increase in cytokine/chemokine signaling, including the IL-1, IFN-γ, IL5, WNT5A, LIF, CXCL12 etc (Supplementary Fig. 10F, quadrant III and bar graph).

### MSH2 binds to IFNAR1 promoter and suppresses its gene expression and signal transduction in BLBC

Since MSH2 and MLH1 work together for MMR, we reason that the distinct function of MSH2 overexpression in BLBC is likely due to other MMR-independent mechanisms. We performed CHIPseq experiments in MDA-MB-231 cells with MSH2-KO or MLH1-KO, reconstituted with HA-tagged MSH2 or MLH1 respectively. We found that MSH2 bound to a total of 1321 regions and MLH1 to 552 regions (Fig. 5A). Among all the MSH2-bound gene loci, *IFNAR1* locus has several pronounced MSH2 peaks, including directly upstream of the promoter area and one in intron 1 (Fig. 5B), all peaks containing either 5 AT repeats or 5 GT repeats (Fig. 5B). MSH2-KO in MDA-MB-231 cells resulted in significant increase in IFNAR1 gene expression, but not IFNAR2 or IFNGR1 (Fig. 5C); and a similar trend for INFAR1 in 4T1 tumors (Fig. 5D), but with significant decrease in INFGR1 by MSH2-KO (Fig. 5D). Datamining the TCGA human breast cancer dataset, we identified a negative correlation between MSH2 protein expression and IFNAR1 gene expression (Fig. 5E); the negative correlation is statistically significant within the BLBC subtype but not the luminal subtype (Fig. 5F). We also determined IFNAR1-mediated signaling transduction in two human cell lines with MSH2 KO. After 3 hrs of serum starvation, IFN-α2 induced significant tyrosine 701 phosphorylation in both WT and KO MDA-MB-231 cells or BT549 cells (Fig. 6A), with significant increase of phos-STAT1/STAT1 – indicative of more efficient IFNAR1 activation – in the KO cells after 1 hr treatment (Fig. 6B, solid, KO; open, WT). We did notice the lower level of STAT1 total protein in the KO cells (Fig. 6A), likely due to an unknown negative feedback from persistent STAT1 activation in the KO cells. It has been shown that IFNAR1 – upon activation by type I interferon, is endocytosed and degraded by lysosome (Constantinescu et al., 1994; Kumar et al., 2003); hence, the persistent IFN signaling transduction requires re-expression of IFNAR1. This is not the case for co-receptor IFNAR2 that is recycled to the plasma member after activation. Since MSH2-KO potentiates the persistent phos-STAT1 signaling, we determined the IFNAR1 protein level after IFN-α2 treatment. As known in the literature, IFN-α2 induced IFNAR1 downregulation in both WT and KO cells (Fig. 6C-D); whereas the kinetics of IFNAR1 degradation and regeneration were very different, with MSH2-KO having faster and stronger IRNAR1 regeneration after 6 or 22 hrs of IFN-α2 treatment (Fig. 6E). Phos-STAT1 levels peaked at 4 hrs and diminished after 6 hrs after IFN-α2 treatment (Fig. 6F). As a major target of type I IFN, CXCL10 expression (Sistigu et al., 2014) was significantly induced by IFN-α2 after 8 hrs and MSH2-KO led to a distinct induction of CXCL10 at the later stages (Fig. 6G), in agreement with the persistent induction of phos-STAT1 signaling (Fig. 6A). To exclude the off target effect of CRISPR/Cas9 during the MSH2 KO, we re-expressed Hibit-tagged MSH2 in the MSH-KO MDA-MB-231 cells (Fig. 6H). After serum starvation and 4 hrs of IFN-α2 treatment, we found that hibit-MSH2 re-expression suppressed the persistent IFN signaling (Fig. 6I). The re-expression experiment also precludes the involvement of mutations due to MSH2-KO.

**Figure 5.**
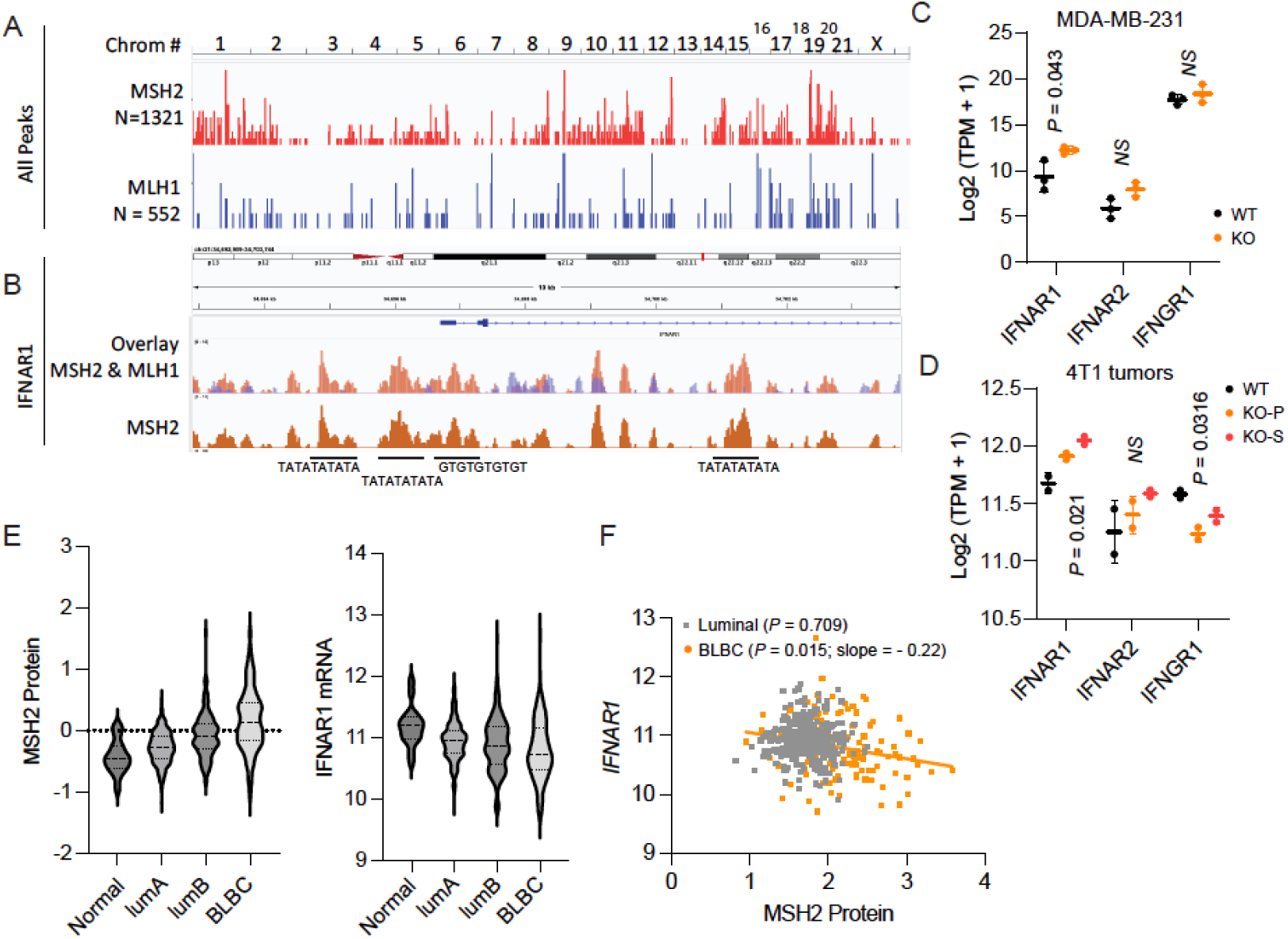
MSH2 binds to *IFNAR1* promoter and represses its gene expression. **A-B.** CHIPseq showing differential DNA binding capacity of MSH2 and MLH1. MDA-MB-231 cells with MSH2-KO or with MLH1-KO were re-expressed HA-tagged MSH2 or MLH1 respectively. Anti-HA antibody was used for CHIP-seq, using anti-HA antibody against nuclear extracts from MSH2-KO or MLH1-KO cells as negative controls. (A) Showing 1321 total peaks for MSH2 and 552 peaks for MLH1; (B) showing specific binding of MSH2 to several loci adjacent to the promoter region of IFNAR1, with unique repeat sequences labelled under the major peaks. **C-D.** MSH2-KO led to increased expression of IFNAR1 gene in (C) human MDA-MB-231 cells and (D) 4T1 tumors based on RNA sequencing of total RNAs from (C) WT or MSH2-KO of MDA-MB-231 cells or (D) from WT or KO 4T1 tumors formed in female Balb/C mice (KO-P, MSH2 KO pooled cells; KO-S, MSH2 KO single clone. **E-F**. MSH2 protein is inversely correlated with IFNAR1 expression in human breast cancers. TCGA breast cancer data were downloaded via UCSC Xena browser and re-analyzed based on breast cancer subtypes (HER2+ excluded due to the relative low numbers). (E) showing MSH2 protein expression (left) versus IFNAR1 gene expression (right) and (F) showing the negative correlation between MSH2 protein and IFNAR1 in BLBC, but not in luminal breast cancers.

**Figure 6.**
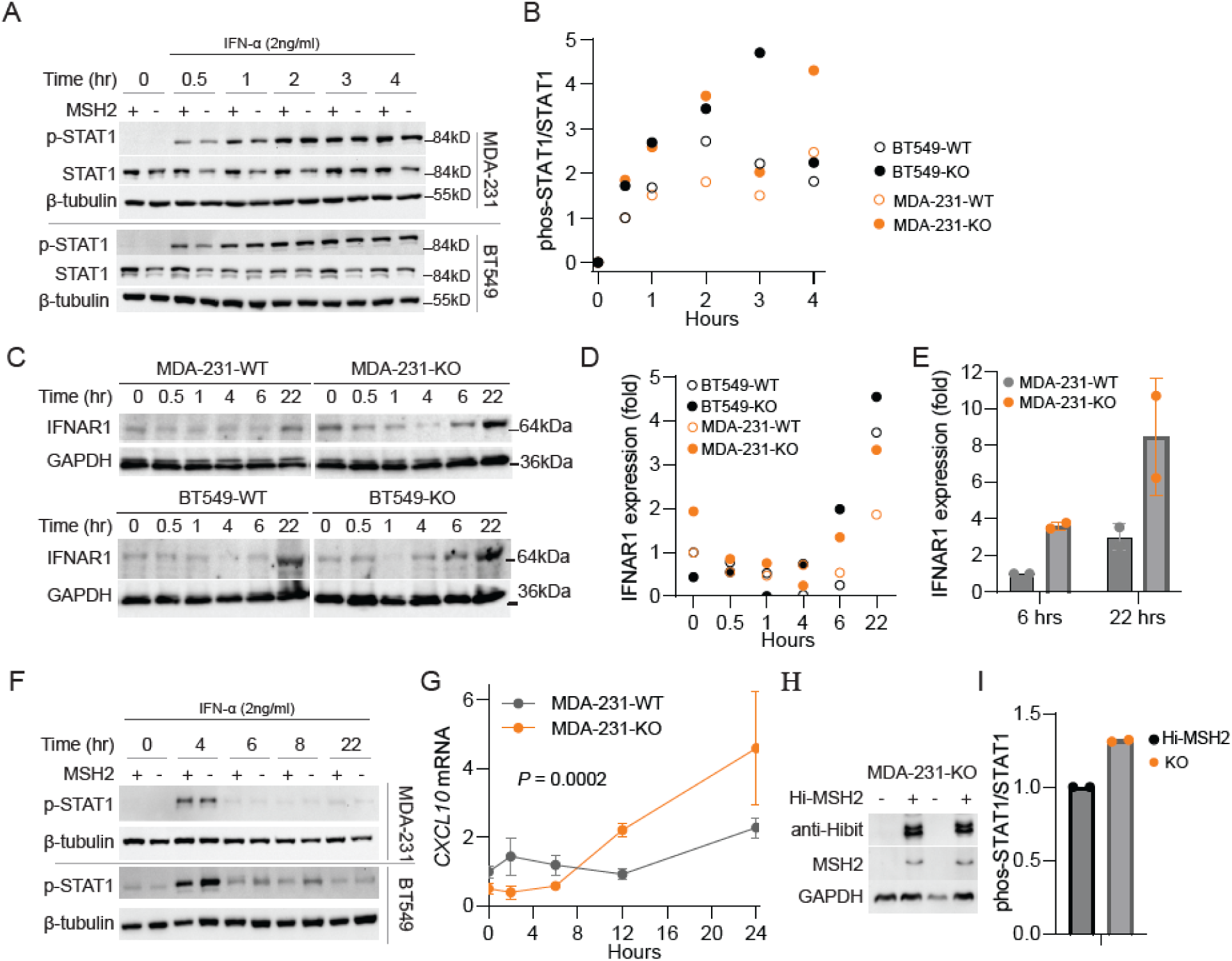
MSH2-KO enhances IFNAR1-mediated signaling transduction by speeding up the IFNAR1 recovery upon persistent interferon stimulation. **A-B.** MSH2-KO leads to persistent signal transduction mediated by IFN/IFNAR1. Cells were serum starved for 3 hrs, following with 2 ng/ml of IFN-α2 treatment of the indicated period. (A) Immunoblotting of phosphorylated STAT1 and STAT1, using tubulin as loading control in WT or KO MDA-MB-231 cells or BT549 cells and (B) Ratios of phos-STAT1/STAT1 based on the densitometry of immunoblots in (A), normalized to tubulin. **C-E.** MSH2-KO leads to faster recovery of IFNAR1 upon persistent IFN-α2 treatment. Cells were serum starved for 3 hrs, following with 2 ng/ml of IFN-α2 treatment of the indicated period. (C) Immunoblotting of IFNAR1, using tubulin as loading control in WT or KO MDA-MB-231 cells or BT549 cells and (D) The IFNAR1 densitometry of immunoblots in (C), normalized to tubulin. (E) IFNAR1 densitometry of 2 independent immunoblots, normalized to GAPDH. F. Impact of MSH2-KO on long term IFNAR1-mediated phos-STAT1, similarly treated as in (A) with extended upto 22 hrs of IFN-α2 treatment. **G.** MSH2-KO leads to elevated *CXCL10* production upon persistent IFN-α2 treatment. WT or KO MDA-MB-231 cells were similarly serum-starved and treated. Real-time PCR was used to determine the expression level of *CXCL10* gene (n=3). **H-I.** Reconstitution of MSH2 suppresses IFNAR1-mediated signaling transduction. MDA-MB-231 KO cells were infected with lentiviral particles encoding Hibit-tagged MSH2 and selected with hygromycin for stable pool culture. (H) Cell lysates from two separated pools were collected and the expression of Hibit-MSH2 was confirmed by using anti-Hibit antibody or anti-MSH2 antibody and (I) KO and hibit-MSH2 expressed cells were serum starved and treated with IFN-α2 for 4 hrs. Ratios of phos-STAT1/STAT1 based on the densitometry of immunoblots were determined and normalized to GAPDH.

Our results support that MSH2 directly binds to the promoter region of IFNAR1 locus and suppresses its expression and that MSH2 KO in BLBC leads to elevated IFNAR1 and potentiates the persistent signaling transduction.

### MSH2 KO leads to exhaustion and sensitizes BLBC to immunotherapy

To study the passage-dependent effect of MSH2 in BLBC, we used LentiCRISPRV2 system to KO MSH2 within MDA-MB-231 cells and culture them up to 30^th^ passages. We confirmed MSH2 KO in these cells even after 30^th^ passage (Supplementary Fig. 11) and performed RNAseq of earlier passages (P5) and late passages (P30) (Fig. 7A). Interestingly, MSH2 KO in the basal B MDA-MB-231 cells led to a passage-dependent shift from an early immune activation at P5 – evidenced by elevation of CD40, CD27, ICOSLG etc – to an exhaustion phenotype supported by elevated expression of immune checkpoints PVR, PDCD1LG2 (PD-L2) and CD274 (PD-L1) (Fig. 7A), in agreement with the human BLBC data showing reverse correlation of MSH2 protein with PVR and CD274 (Supplementary Fig. 10C). We also determined CD274 expression in 4T1 cells and found consistent data that MSH2-KO led to 7.7 fold increase in CD274 expression and several chemokines, but not in the MLH1-KO cells (Fig. 7B). Using the 4T1 BLBC model, we established a cell lysate vaccine model to boost the immune activation within 4T1 tumors (Fig. 7C), otherwise known as an unresponsive tumor model to immune checkpoint inhibitors (Sagiv-Barfi et al., 2015). We found that after vaccination, *Msh2*-KO led to an improved survival (Fig. 7D) and reduced tumor growth (Fig. 7E), suggesting vaccination sensitizes *Msh2*-KO effect even in the primary tumors. Vaccination + anti-PD-1 improved survival (Fig. 7D) and inhibited tumor growth (Fig. 7E) in the WT 4T1 tumor bearing mice, but further extended survival and reduced tumor growth in *Msh2*-KO tumor bearing mice (Fig. 7D-7E). Flow cytometric analysis supported that *Msh2*-KO led to increased PD-1^+^TIM-3^+^ exhausted T cells and anti-PD-1 treatment reduced that population (Fig. 7F-7G).

**Figure 7.**
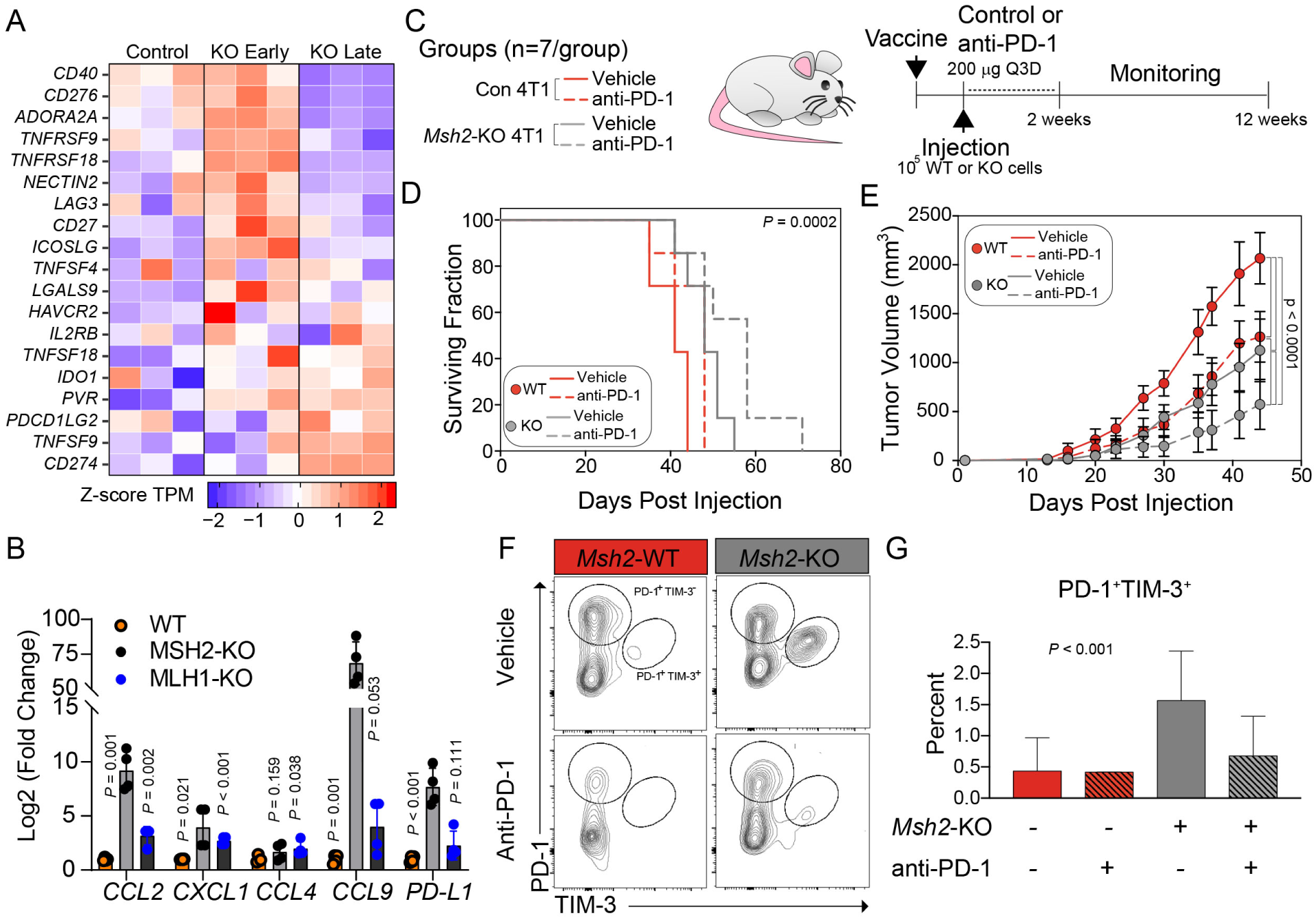
Long-term MSH2 KO induces an exhaustion phenotype of T cells and sensitizes tumors to anti-PD-1 therapy. **A.** MSH2 KO increases the expression of immune checkpoints in MDA-MB-231 BLBC cells in late passages (P30). Lentiviral particles encoding Cas9 and guide RNA for MSH2 were used to infect MDA-MB-231 cells, following 5-day selection process with puromycin to induce MSH2 KO. The MSH2 KO cells were passaged to P5 or P30, following with RNAseq and differential expression genes (DEGs). Control cells were infected lentiviral particles encoding Cas9 and scramble control. n = 3 per group. **B.** Real-time PCR showing the expression of chemokines or CD274 (PD-L1) in control, MSH2-KO, or MLH1-KO 4T1 cells (n=4). **C-E**. Long-term MSH2 KO in 4T1 model leads to T cell exhaustion and sensitizes tumors to anti-PD-1 therapy. (C) Experimental diagram, including whole 4T1 cell vaccine 7 days before tumor cell injection. Anti-PD-1 treatment starts one day after tumor cell injection, every 4 days upto 2 weeks. (D) survival curves and (E) tumor growth curves were shown (n = 7 per group). **F-G**. MSH2 KO induces T cell exhaustion and sensitizes tumors to anti-PD-1 treatment. (F) Example of flow cytometry using tumors in D, using PD-1+TIM-3+ as exhaustion T cell population; and (G) Histogram showing the statistics of PD-1+TIM-3+ T cell population (n=4-5 per group).

## Discussion

Known as a tumor suppressive pathway, MMR genes are mostly considered equal in relation to cancer initiation, hypermutator phenotype, high level of mutational load and the sensitivity to ICIs. Based on several important trials (Geoerger et al., 2020; Le et al., 2020; Le et al., 2015; Marabelle et al., 2020), FDA approved pembrolizumab for first-line treatment of patients with microsatellite instability-high (MSI-H) or mismatch repair deficient (dMMR) cancers, irrelevant of cancer types. Interestingly, we found that *MSH2* expression is frequently elevated in human BLBC and several other cancer types and its protein expression is correlated with worse prognosis in human BLBC. A published study based on immunohistochemistry agrees with our finding that MSH2 protein is higher, while MLH1 protein is lower, in BLBCs than other breast cancer subtypes (Alkam et al., 2013). Mouse models also support the tumor/metastasis-promoting function of MSH2, in contrast to the role of MLH1 in tumor/metastasis suppression. This is the first report showing opposite functions of MMR genes in metastasis.

Despite the established role of MSH2 as a tumor suppressor (Bonadona et al., 2011; Fishel et al., 1993; Papadopoulos et al., 1994; Reitmair et al., 1996), a recent proteomic screen for synthetic lethality demonstrated mismatch repair pathway as one of the most sensitive DNA repair pathways for synthetic lethality (Pearl et al., 2015). This may be due to the interaction of mismatch repair proteins with multiple DNA repair pathways including homologous recombination (Spies and Fishel, 2015) that is defective in BRCA1-mutated BLBC. If genomic instability can be thought of as a delicate balance between clonal evolution of cancer cells and cell death, the interaction of MMR machinery with multiple DNA repair pathways makes it an ideal target for therapeutic study in BLBC. We did observe that MSH2 protein expression is correlated with less in frame deletion (in-del) mutations in the BLBC (data not shown), supporting the expected role of MSH2 in DNA repair. The surprises come from the observation of differential immune regulations from MSH2- or MLH1-deletion from tumor cells, where MSH2-deletion leads to the collective expression of chemokines and cytokines, as well as exhaustion markers for T cells in both mouse and human BLBCs. The chemokine induction is apparently the primary cause of immune cell infiltrations within the TME, associated with T cell activation and exhaustion phenotypes. MSH2 promotes metastasis also relies on immune cells since the MSH2-mediated metastasis is abolished in the NSG mice.

At the mechanistic level, we found that MSH2 is directly involved in the expression of IFNAR1 – the receptor subunit for a master immune regulation program initiated by type I IFN. MSH2 binds to several genetic elements upstream or downstream of IFNAR1 promoter area; all of the genetic locations consist of 5 AT or GC repeats. The binding of MSH2 with those genetic elements is correlated with decreased IFNAR1 expression as MSH2-KO leads to increased IFNAR1 and a faster protein recovery after persistent IFN-α2 treatment, which leads perpetuate and enhanced expression of IFN target gene such as CXCL10. Other IFNAR1-downstream genes include immune checkpoints such as PD-L1 and PVR that are elevated in the MSH2-KO BLBC cells. Indeed, MSH2-KO sensitizes tumors to immune checkpoint inhibitors in mouse models of breast cancer.

MLH1 consistently behaves as a tumor/metastasis-suppressor whose deletion induces elevated tumor growth as well as increased metastasis. MLH1 KO did not lead to significant increase in TI-immune cells, which is a bit surprising considering that the MLH1 KO cells were derived from the combination of several clones derived from a single cell. The single clones underwent at least > 23-24 doubling times before mixed and injected into the mice (Fig. 2), which had been expected to generate sufficient neoantigens for immune activation and immune cell infiltration. The human BLBC dataset agrees with the irrelevance of MLH1 expression with immune infiltration because MLH1 expression levels are not correlated with any immune modulatory pathways; rather, MLH1-low BLBCs exhibit active ribosomal and ribosomal activities, supporting active protein synthesis and metabolic functions (Supplementary Fig. 9) – likely pointing to the aggressive behavior of those BLBCs.

The general increase of MSH2 expression in some other cancer types, either by amplification or increased expression (Supplementary Fig. 2), points to a potentially common mechanism for MSH2 to promote tumor progression. At this stage, it is too early to know whether MSH2 or MutSα is a potential therapeutic target for cancer therapy. It is tempting to inhibit MutSα within the BLBC or other cancer types for potential therapeutic development, which is expected to suppress cancer progression and at the same time to sensitize cancers to immune checkpoint inhibitors.

## Methods

### Reverse-phase Protein Analysis

RPPA data was downloaded from the TCPA Portal located at http://tcpaportal.org/. Data was processed as previously described and clinical data was attached by Patient/Sample ID (Borcherding et al., 2018). Protein-based survival analysis was performed using the previously-developed TRGAted R Shiny application, code and processed data for all TCGA cohorts are located at https://github.com/ncborcherding/TRGAted. (Borcherding et al., 2018) Molecular-subtype designation was based on the TCGA Analysis Working Group pipeline, fitting PAM50 subtypes based on RNA-seq data. Protein versus mRNA correlations was performed using log2(x+1) mRNA quantification downloaded from the cBioPortal (Cerami et al., 2012; Gao et al., 2013). Correlations for both protein-mRNA and protein-protein comparisons utilized the rank-based Spearman approach.

### RNAseq

Total RNAs were purified directly either from cell lines (WT or MSH2-KO of BT549 or MDA-MB-231 cells) or tumor specimens (WT, MSH2-KO or MLH1-KO of Py8119 or 4T1 tumors). Illumina Standard library construction (New England Biolabs) and sequencing (Illumina NovaSeq, S4, 2×150bp) were performed at the ICBR core facility, the University of Florida Health Cancer Center (UFHCC). RNA-seq fastq files were trimmed using seqtk and the quality assessed using FastQC. The trimmed reads were aligned to the mouse genome (mm10) or human genome (hg38) using Hisat2 and transcript counts were obtained using HTseq. DeSeq2 was used for differential expression analysis (Lorentsen et al., 2018). Significance for RNA-seq data was determined using DeSeq2 (Love et al., 2014).

### Differential Gene and Pathway Analysis

*G*ene-level HTSeq count data was downloaded from the UCSC Xena Browser at http://xena.ucsc.edu/. Using the DESeq2 R package (v1.16.1), count data were separated in half by MSH2 RPPA quantification and converted into negative binomial distributions. A parametric model was fitted for the data and significance and log2 fold-change was computed using Wald testing. P-values for the differential analysis was corrected for multiple hypothesis testing using the Benjamini and Hochberg method. Significance threshold was set at adjust P-value ≤ 0.05 and log2-fold change ≥ |0.5|. A total of 2,094 genes met the criteria, with 650 genes significantly increased in MSH2-high BLBC samples and 1,444 genes significantly decreased. For visualization of the expression of specific genes by MSH2 grouping, the expression matrix was regularized log transformed. Ingenuity Pathway Analysis was performed using the above significance cut-off. DESeq-based log2-fold change was incorporated with the bias-corrected Z-score for upstream regulator analysis in order to visualized activation relative to mRNA fold-change.

### RNA-immune cell estimates

Immune and stromal ESTIMATE scores for all breast cancer samples in the TCGA were downloaded (http://bioinformatics.mdanderson.org/estimate/). BLBC samples with MSH2 protein quantifications were extracted and split by mean MSH2 value. In addition to general immune cell difference, log2 RSEM RNA values for BLBC samples were imported into the CIBERSORT portal, in order to estimate the contribution of individual immune lineages. Absolute estimates of immune cell populations were combined for total immune cell lineages and then were then divided by mean MSH2 protein values.

### Cell lines and cell culture

MC38 mouse colon carcinoma cells (Kerafast Inc., Boston, MA) were maintained in DMEM supplemented with 10% FBS, 1mM glutamine, 0.1M non-essential amino acids (Thermo Fisher, Waltham, MA), 1 mM sodium pyruvate, 10 mM HEPES, 100 U/ml penicillin and 100 µg/ml streptomycin. 4T1 mouse mammary carcinoma cells were purchased from American Type Culture Collection (ATCC) and cultured in RPMI medium supplemented with 10% fetal bovine serum (FBS) 100 U/ml penicillin and 100 µg/ml streptomycin. MDA-MB-231 and BT549 human breast cancer cell lines and Py8119 mouse mammary carcinoma cells were purchased from ATCC and cultured in DMEM supplemented with 10% fetal bovine serum (FBS), 100 U/ml penicillin and 100 µg/ml streptomycin. All cells were cultured at 37 °C and 5% CO2 in a humidified incubator.

### CRISPR/CAS9-mediated knockout and viability assay

The knockout clones of MC38, Py8119 and 4T1 cells were generated by transient transfection of Cas9 protein-guide RNA complex into parental cells, followed by single clone section and immunoblot to confirm KO of MSH2 or MLH1 following the instruction (Synthego). The knockout clones MDA-MB-231 and BT549 cells were constructed either by transient transfection of Cas9 protein-guide RNA complex or by lentiviral particles co-expressing Cas9 and guide RNA (LentiCRISPRv2)(Stringer et al., 2019), following by puromycin selection for stable pooled culture. Cells were passages every 3 days and passage numbers were recorded.

Human MSH2 CRISPR guide #1 sequence (G1):

5’ - CACCGCTTCTATACGGCGCACGGCG - 3’

3’ - CGAAGATATGCCGCGTGCCGCCAAA - 5’

Human MSH2 CRISPR guide #2 sequence (G2):

5’ - CACCGGCGCCGTATAGAAGTCGCCC - 3’

3’ - CCGCGGCATATCTTCAGCGGGCAAA - 5’

Mice MSH2 CRISPR guide sequence:

5’ - CACCGGCGCCGTGTAAAAGTCGCCG - 3’

3’ - CCGCGGCACATTTTCAGCGGCCAAA - 5’

DNA oligonucleotides for gRNA sequences and their reverse complement sequences plus adapters were annealed, ligated into lentiCRISPRv2 vector for lentivirus packaging in HEK293T cells and infection and puromycin selection of different cells used in this study.

For RNA & Protein complex (RNP)-based MSH2 or MLH1 KO used in Figure 2 and Supplementary Figures 3-5, guide RNAs using the above sequences and Cas9 protein were purchased (Synthego). RNP complex was transiently transfected into cells per company-provided instructions and single clones were selected for culture and expansion. KO of target proteins were validated using immunoblotting.

### Animal studies

Animal studies were approved by the University of Florida Institutional Animal Care and Use Committee (IACUC: 202110399). Animals were housed in a pathogen-free facility accredited by the Association for Assessment and Accreditation of Laboratory Animal Care at the University of Florida. Temperature is 21.1-23.3 °C with daily fluctuation of 2°C. The humidity is 50%, but can vary +/-15% daily. The photoperiod is 12:12 and the light intensity range is 15.6-25.8 FC. C57BL/6 and Balb/C mice were purchased from Charles River Laboratories (San Diego, CA). NSG mice were purchased from Jackson Laboratories (Bar Harbor, ME). For syngeneic tumor models, 500 cells (for Py8119) were resuspended in 50% phosphate buffered saline (PBS) + 50% Matrigel (Corning Inc., Corning, NY) and implanted into 6-8 week old C57BL/6 (MC38, *s.c*; and Py8119, intra mammary fatpad), Balb/C (4T1, intra mammary fatpad), or NSG mice (MDA-MB-231, intra mammary fatpad). Tumor growth was monitored daily and measured twice weekly with calipers and volume was determined using the formula ½ (L x W2).

### Flow cytometry

Flow cytometry procedure is adapted from our recent publications (Kolb et al., 2021). Briefly, 200 mg of tumor tissue was enzymatically and mechanically digested using the mouse Tumor Dissociation Kit (Miltenyi Biotec). For blood and spleens, red blood cells were lysed using ACK lysis buffer and mononuclear cells were isolated by density gradient using SepMate Tubes (StemCell Technologies) and Lymphoprep density gradient media (StemCell Technologies). Mouse cells were then washed and incubated with combinations of the following antibodies: anti-mouse CD62L-BV785 (clone MEL-14), anti-mouse MHCII I-A/I-E-BB515 (Clone 2G9, BD Biosciences), anti-mouse CD11B-PEdazzle (clone M1/70), anti-mouse CD45-AF532 (clone 30F.11), anti-mouse CD3-APC (clone 17A2), anti-mouse CD8-BV510 (clone 53-6.7), anti-mouse CD4-BV605 (clone GK1.5), anti-mouse NK1.1-AF700 (clone PK136), anti-mouse CD69-SB436 (clone H1.2F3, eBioscience), anti-mouse CD279 (clone-PerCP-EF710, eBioscience Inc), anti-mouse CD366-PacBlue (clone B8.2c12), anti-mouse CD11C-PE-Cy7 (clone N418), anti-mouse Ly6G-FITC (clone IA8), anti-mouse Ly6C-BV711 (clone HK1.4) anti-mouse F4/80-BV650 (clone BM8), anti-mouse CD80-BV480 (clone 16-10A1, BD Biosciences), anti-mouse CD25-PE-Cy5 (clone PC61) plus FVD-eFluor-780 (eBioscience) and mouse FcR blocker (anti-mouse CD16/CD32, clone 2.4G2, BD Biosciences). After surface staining, cells were fixed and permeabilized using the FOXP3/Transcription Factor Staining Buffer Set (eBioscience). Cells were stained with a combination of the following antibodies: anti-mouse FOXP3-APC (clone FJK-16S, eBioscience), anti-mouse Granzyme B-Pacific Blue (clone GB11), anti-mouse Perforin-PE (clone S16009B), anti-mouse Ki-67-PerCP-Cy5.5 (clone 16A8). Flow cytometry was performed on a 3 laser Cytek Aurora Cytometer (Cytek Biosciences, Fremont, CA) and analyzed using FlowJo software (BD Biosciences). All antibodies are from Biolegend unless otherwise specified.

### Immunoblotting

Cells or tumors samples were lysed with RIPA buffer (150 mM NaCl, 5 mM EDTA, 50 mM Tris pH8.0, 1% sodium deoxycholate, 1% NP-40, 0.5% SDS) supplemented with 1mM dithiothreitol and protease inhibitors. Cell lysates were then separated by SDS-PAGE and analyzed by standard western blotting protocol as we previously published(Kolb et al., 2021).

### Statistical Analysis

Two sample statistical testing principally utilized Welch’s t-test, allowing for unequal variance and sample size, unless otherwise indicated. Multiple comparisons across three or more groups utilized ANOVA with the Tukey honest significant difference adjustment for multiple comparisons.

## Author Contributions

Funding: W.Z. and M.S.;

Conception and design: W.Z., M.S., J.M., N.B., S.J.;

Development of methodology: W.Z., M.S., J.M., N.B., S.J., T.I.T., E.C., K.E.C., M.H., L.W., K.K.A., R.K.;

Acquisition of data: W.Z., M.S., J.M., N.B., S.J., T.I.T., E.C., K.E.C., M.H., L.W., K.K.A., R.K.;

Analysis and interpretation of data: M.S., J.M., N.B., S.J., W.Z.;

Writing, review, and/or revision of the manuscript: M.S., W.Z., R.W., R.K., M.J.;

Supervision: W.Z.

## Acknowledgements

The work was mainly supported by DOD/CDMRP grant BC180227 (W.Z.) and BC180227P (M.S.), and partially supported by NIH grants CA269661 (W.Z.), CA200792 (W.Z.), CA260239 (W.Z.), and DOD/CDMRP grant BC200100 (W.Z.). W.Z. was also supported by an endowment fund from the Dr. and Mrs. James Robert Spenser Family.

## Supplementary information

**Supplementary Table 1. The correlation between patient prognosis and DNA repair genes/proteins in the TCGA BLBC dataset**

Hazard Ratios and –log10(p-value) Variables of Major DNA Repair Genes or Proteins (MSH2 and MSH6) from the TCGA and TPGA Datasets. Greater than 1.3 of −log10(p-value) is considered significant.

**Supplementary Table 2.** Inverse correlation of MSH2 protein with immune modulatory molecules within the TCGA Breast Cancer BLBC dataset (n = 121).

**Supplementary Table 3. Positive** correlation of *MLH1* expression with immune modulatory molecules within the TCGA Breast Cancer BLBC dataset (n = 121).

**Supplementary Table 4. MSH2-low human BLBC specimens have significantly enriched signaling pathways involving in immune regulations**

GSEA analysis using the C5: ontology gene sets comparing MSH2 protein-low versus -high human TCGA BLBC specimens (lowest 1/3 versus highest 1/3), showing positively enriched C5: ontology gene sets in the MSH2-low specimens.

**Supplementary Table 5. MSH2-high BLBC specimens have elevated pathways related to cancer cell proliferation and progression**

GSEA analysis using the C5: ontology gene sets comparing MSH2 protein-low versus -high human TCGA BLBC specimens (lowest 1/3 versus highest 1/3), showing negatively enriched C5: ontology gene sets in the MSH2-low specimens.

**Supplementary Table 6. *MLH1*-low human BLBC specimens have significantly enriched signaling pathways related to active and aggressive cancer types**

GSEA analysis using the C5: ontology gene sets comparing *MLH1* mRNA-low versus -high human TCGA BLBC specimens (lowest 1/3 versus highest 1/3), showing positively enriched C5: ontology gene sets in the *MLH1*-low specimens.

**Supplementary Table 7. *MLH1*-high human BLBC specimens have no significantly enriched signaling pathways related to cancer progression**

GSEA analysis using the C5: ontology gene sets comparing *MLH1* mRNA-low versus -high human TCGA BLBC specimens (lowest 1/3 versus highest 1/3), showing negatively enriched C5: ontology gene sets in the *MLH1*-low specimens.

**Figure S1.**
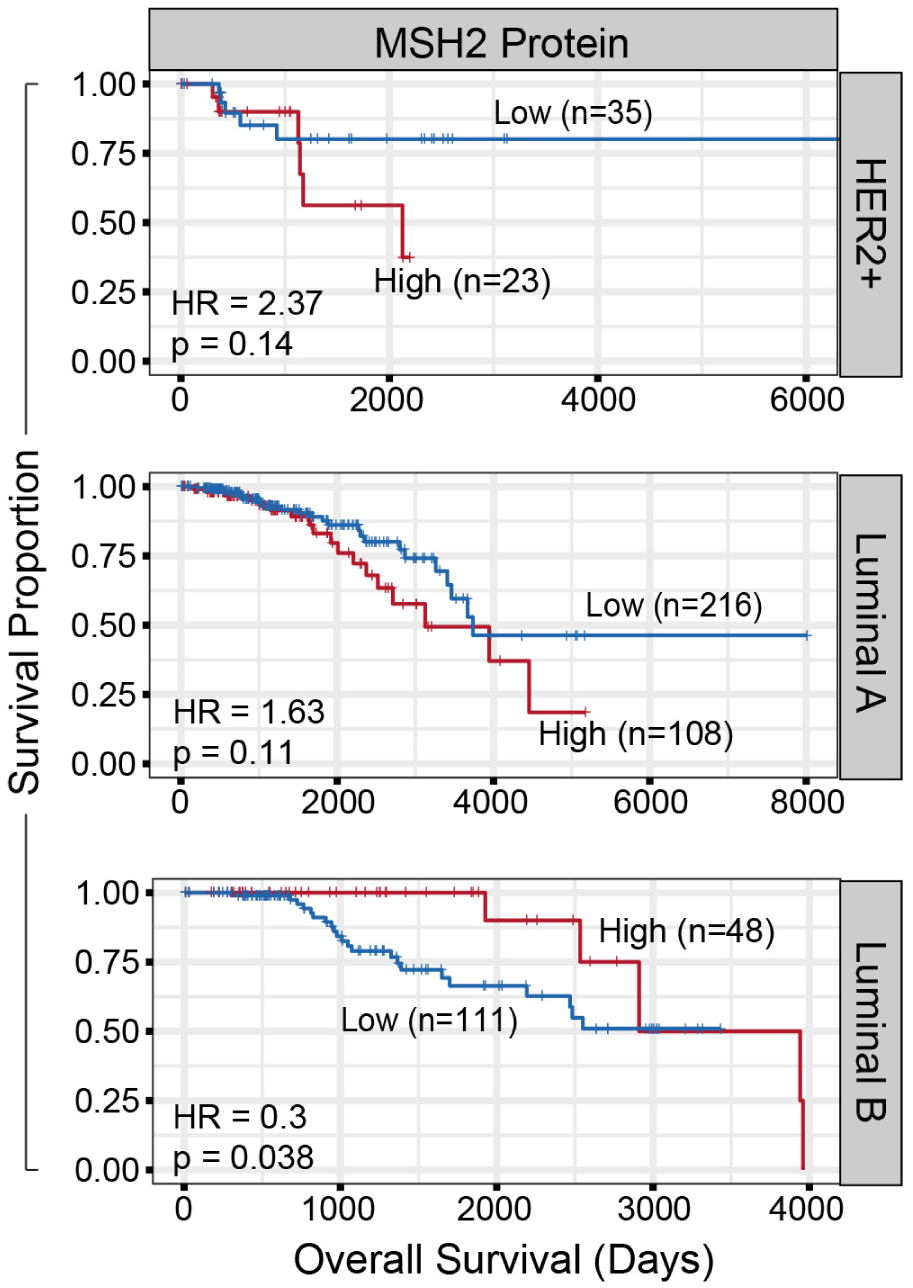
Survival curve based on MSH2 protein levels in the TCGA HER2+, Luminal A, or Luminal B samples, comparing high (red) to low (black). HR and P values are indicated.

**Figure S2.**
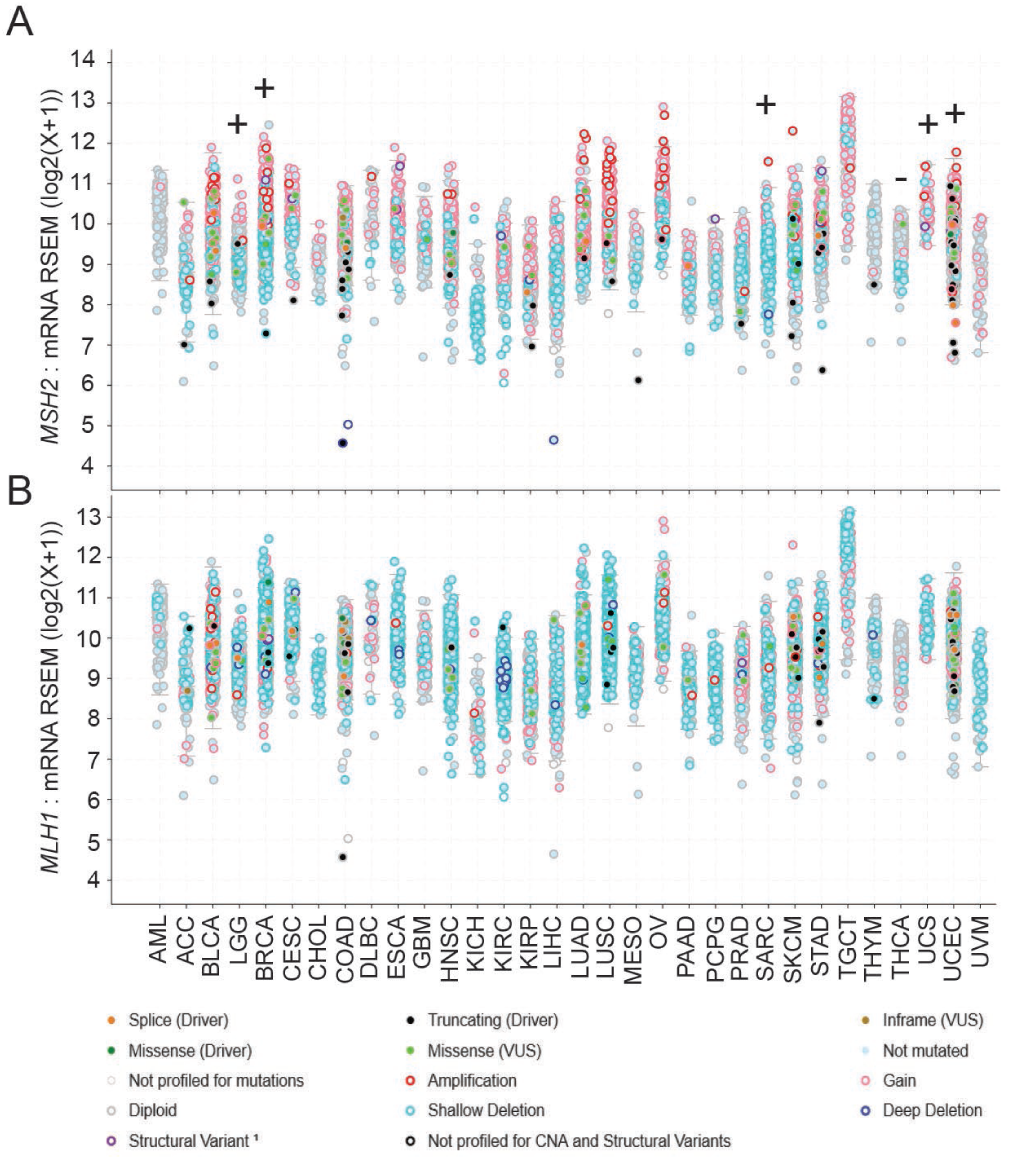
MSH2 – but not MLH1 – exhibits frequent amplification or gain of expression in human cancers. A-B. cBioPortal for Cancer Genomics was used to extract **(A)** *MSH2* or **(B)** *MLH1* mRNA expression and genomic variations from the TCGA PanCancer Atlas Studies (n = 10967). Cancer types are using the standard TCGA abbreviations.

**Figure S3.**
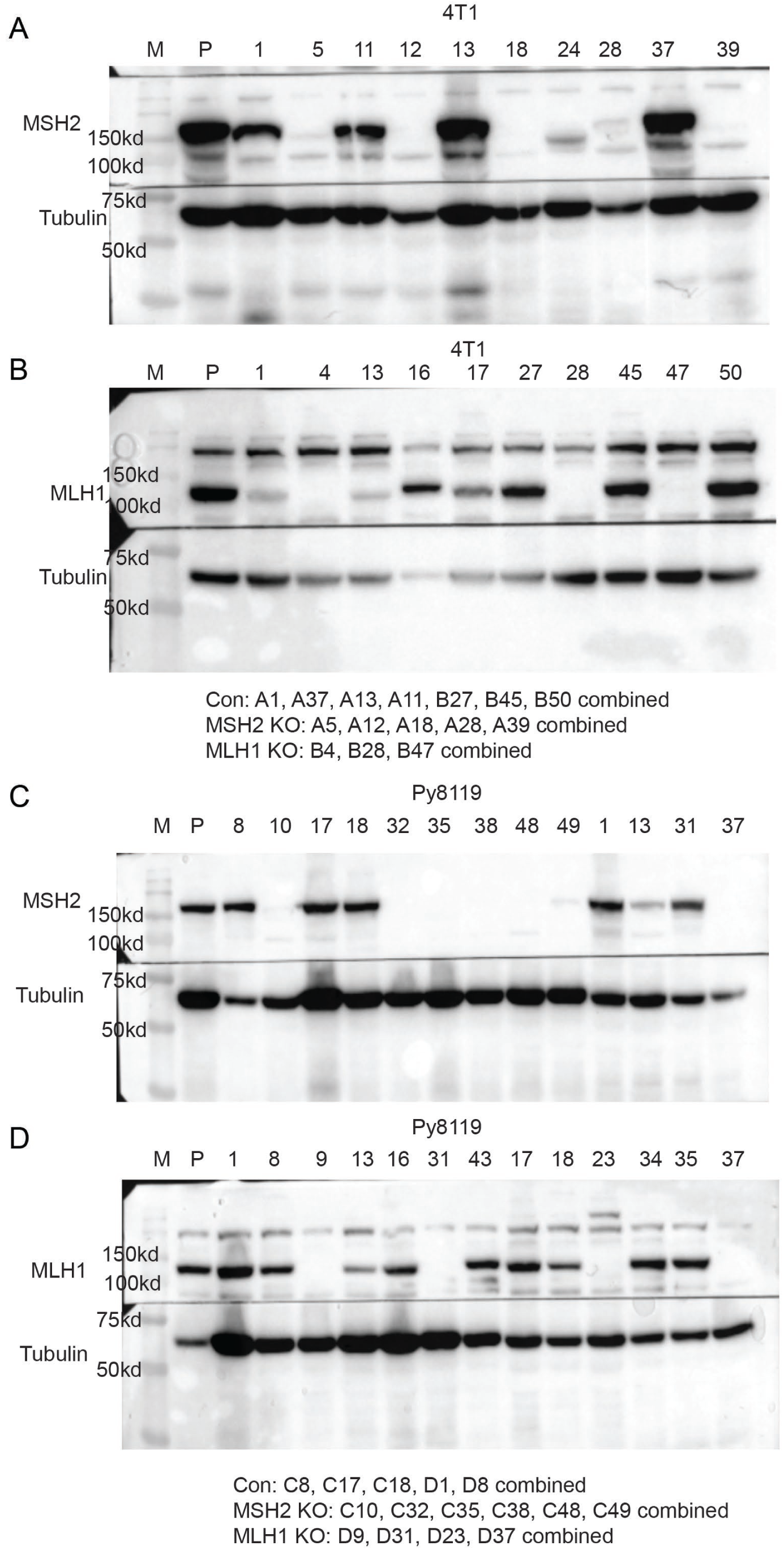
The Generation of MSH2-KO or MLH1-KO cells of Py8119 or 4T1 mouse breast cancer cells using CRISPR/Cas9-based genomic editing method. A-D. Cas9/Guide RNA complex were transiently transfected into Py8119 or 4T1 cells, following with single clone selection, validation using immunoblotting. Several clones were combined for experiments in Figure 2.

**Figure S4.**
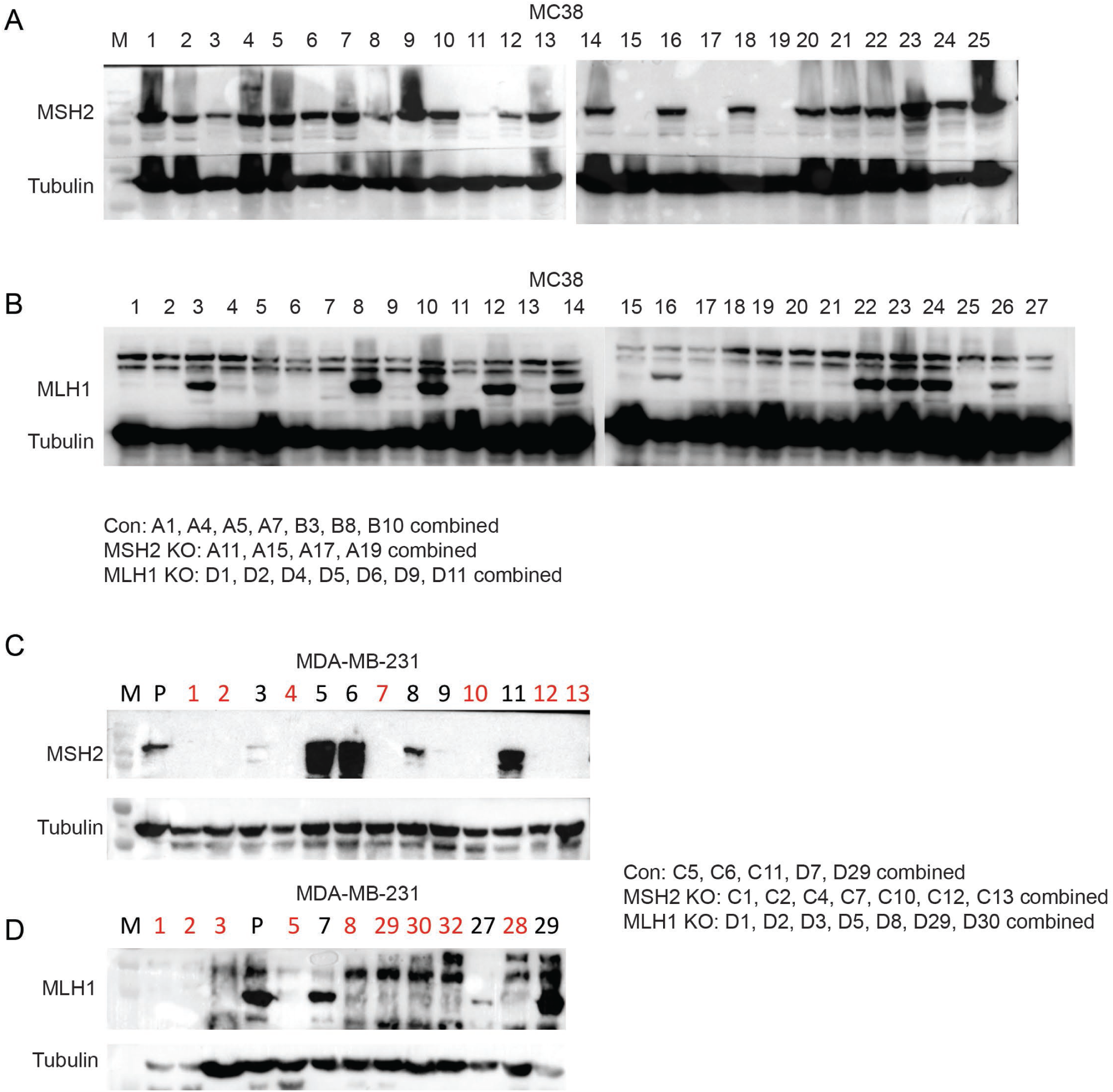
The Generation of MSH2-KO or MLH1-KO cells of MC38 mouse colon cancer cells or MDA-MB-231 human breast cancer cells using CRISPR/Cas9-based genomic editing method. A-D. Cas9/Guide RNA complex were transiently transfected into MC38 or MBA-MD-231 cells, following with single clone selection, validation using immunoblotting. Several clones were combined for experiments in Figure S5.

**Figure S5.**
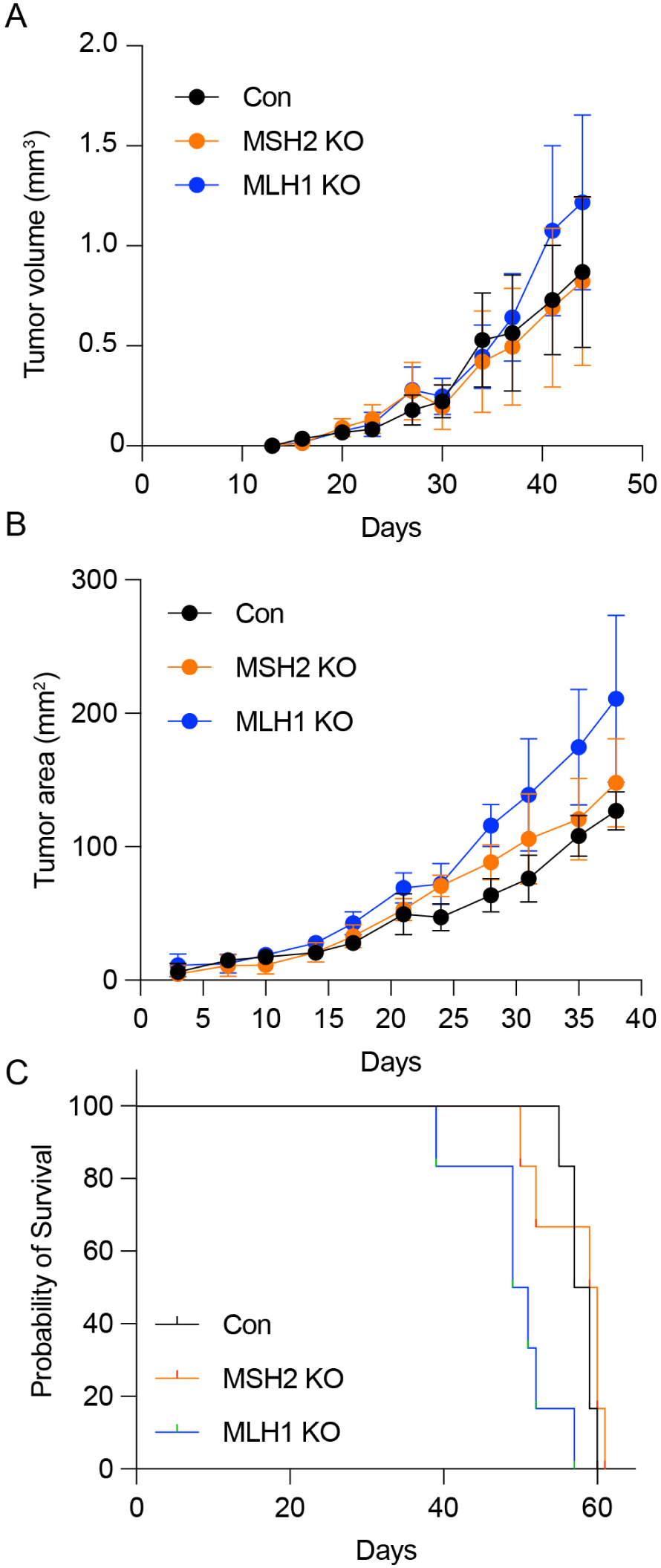
Opposing functions of MSH2 and MLH1 in breast cancer metastasis in immune competent mouse. **A-B.** Primary tumor growth curve (A-B) or survival (C) of control (Con), MSH2-KO or MLH1-KO cells of MC38 cells i.v. injected into 8-week old female C57BL/6J (n = 7 per group) (A) or into #4 mammary fatpad of 8-week old NSG mice (n = 7 per group) (B-C).

**Figure S6.**
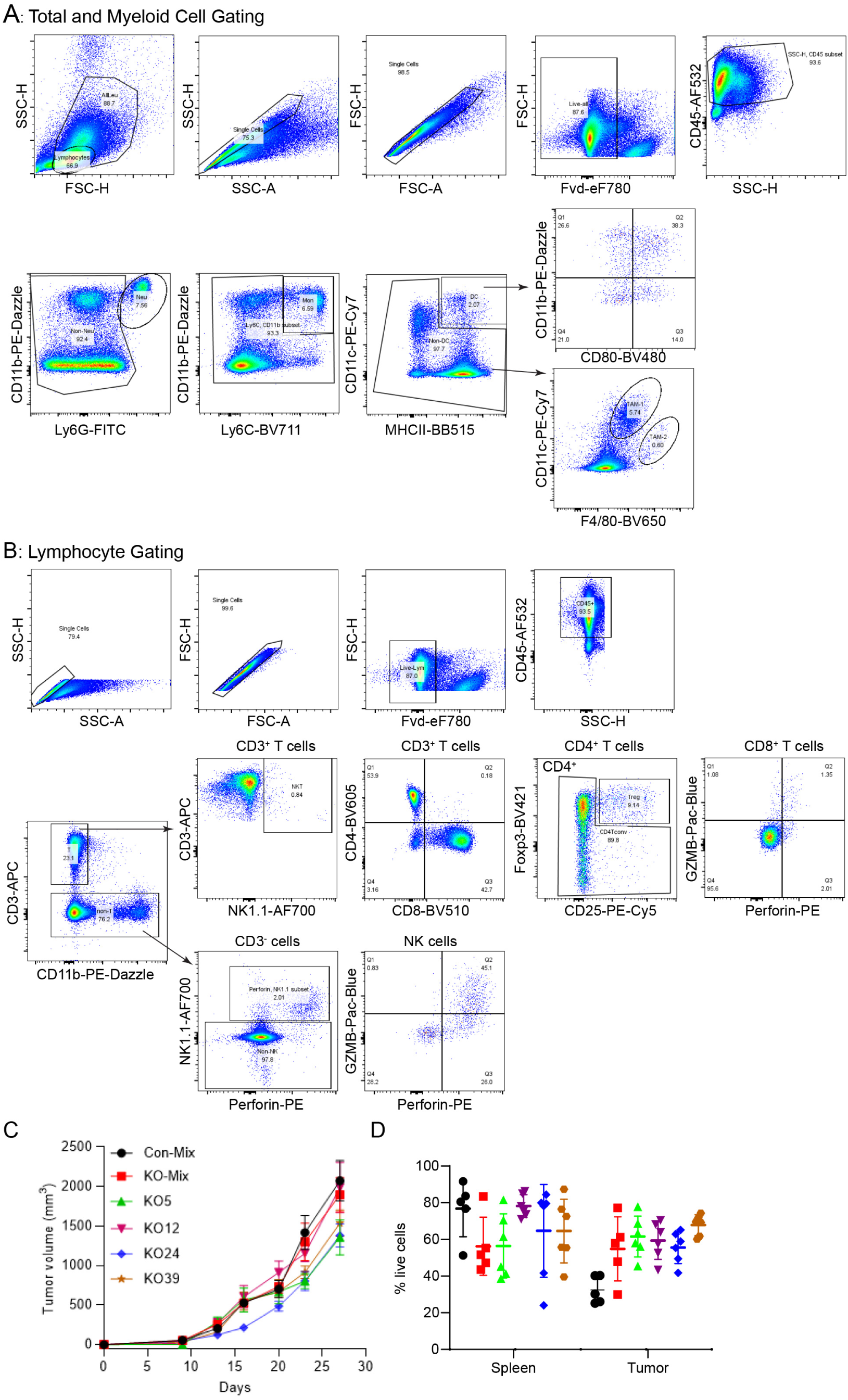
Gating scheme and supplementary information for. Figure 3**. A.** Scheme showing the total live cell and all leukocyte gating. **B.** Scheme showing gating on lymphocytes. **C-D**. Primary tumor growth curve **(C)** or total immune cell infiltrates **(D)** of control-mix clones (Con), MSH2-KO mix clones, or MSH2-KO individual clones of 4T1 orthotopically injected into #4 fatpad of 8-week old female Balb/C mice (n = 7 per group). **(D)** *P* < 0.01 comparing all MSH2-KO groups with Con-mix in total immune cell infiltrations.

**Figure S7.**
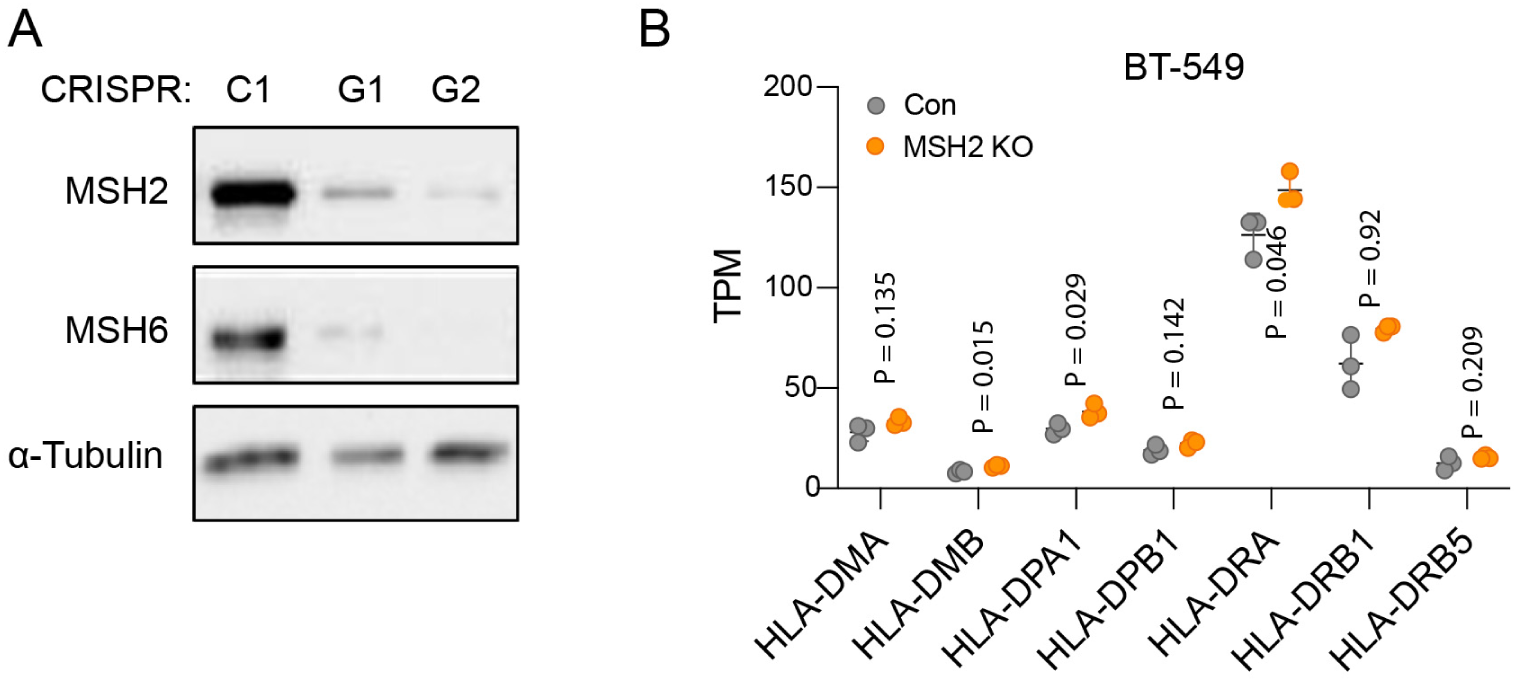
MSH2 KO increases the expression of MHC genes in BT-549 BLBC cells. **A.** Lentiviral particles encoding Cas9 and guide RNA for MSH2 were used to infect BT-549 cells, following 5-day selection process with puromycin to induce MSH2 KO. The MSH2 KO cells were passaged to P30 and KO efficiency was confirmed using immunoblotting, with concomitant loss of MSH6. **B.** RNAseq and differential expression genes (DEGs) were analyzed from WT and MSH2 KO (G2) cells, showing elevation of MHC genes in the MSH2 KO cells.

**Figure S8.**
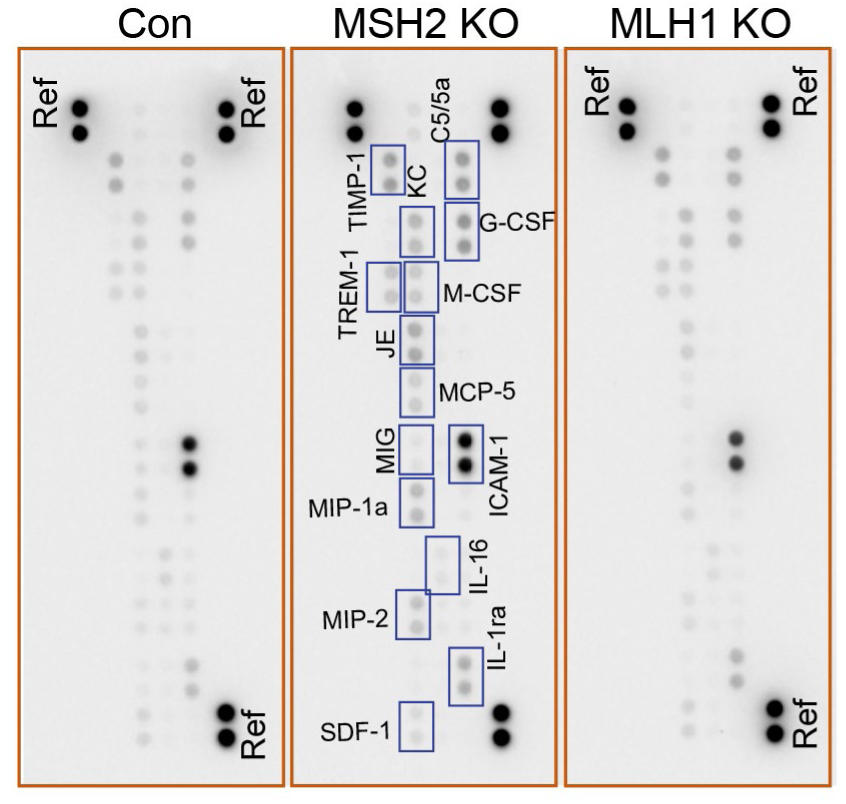
Original images used in and Supplementary information for Fig. 4E (Proteome Profiler Mouse Cytokine Array Kit, Panel A, Biotechne, R&D).

**Figure S9.**
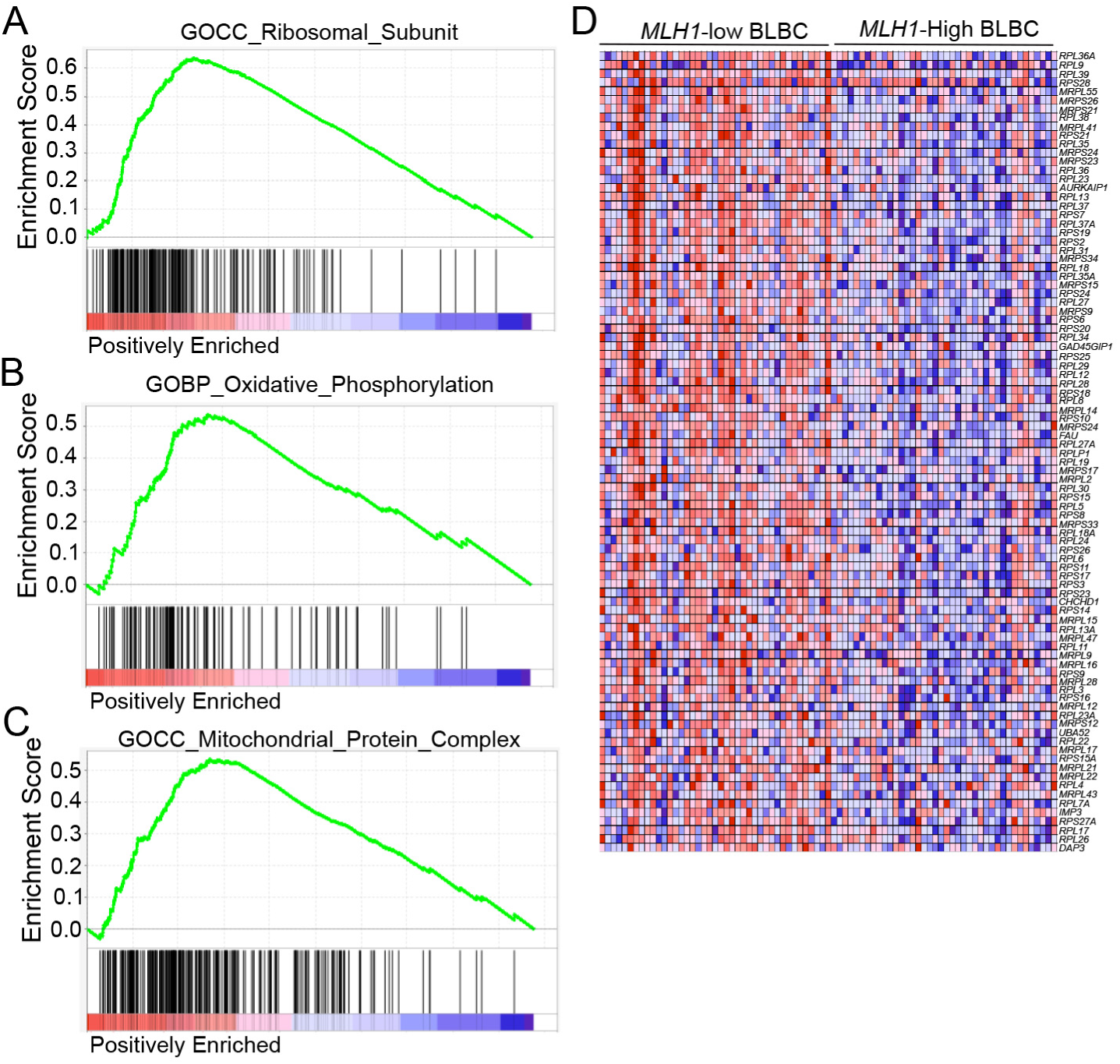
MLH1-low human BLBC specimens have significantly enriched signaling pathways related to metabolism and protein synthesis. Also refer to Supplementary Table 6. GSEA analysis using the C5: ontology gene sets comparing MLH1 mRNA-low versus -high human TCGA BLBC specimens (lowest 1/3 versus highest 1/3), showing positively enriched C5: ontology gene sets in the MLH1-low specimens, including (A) Enriched Ribosomal Subunit; (B) Increased Oxidative Phosphorylation; (C) Mitochondrial Protein Complex; and (D) individual Ribosomal Subunit listed from (A).

**Figure S10.**
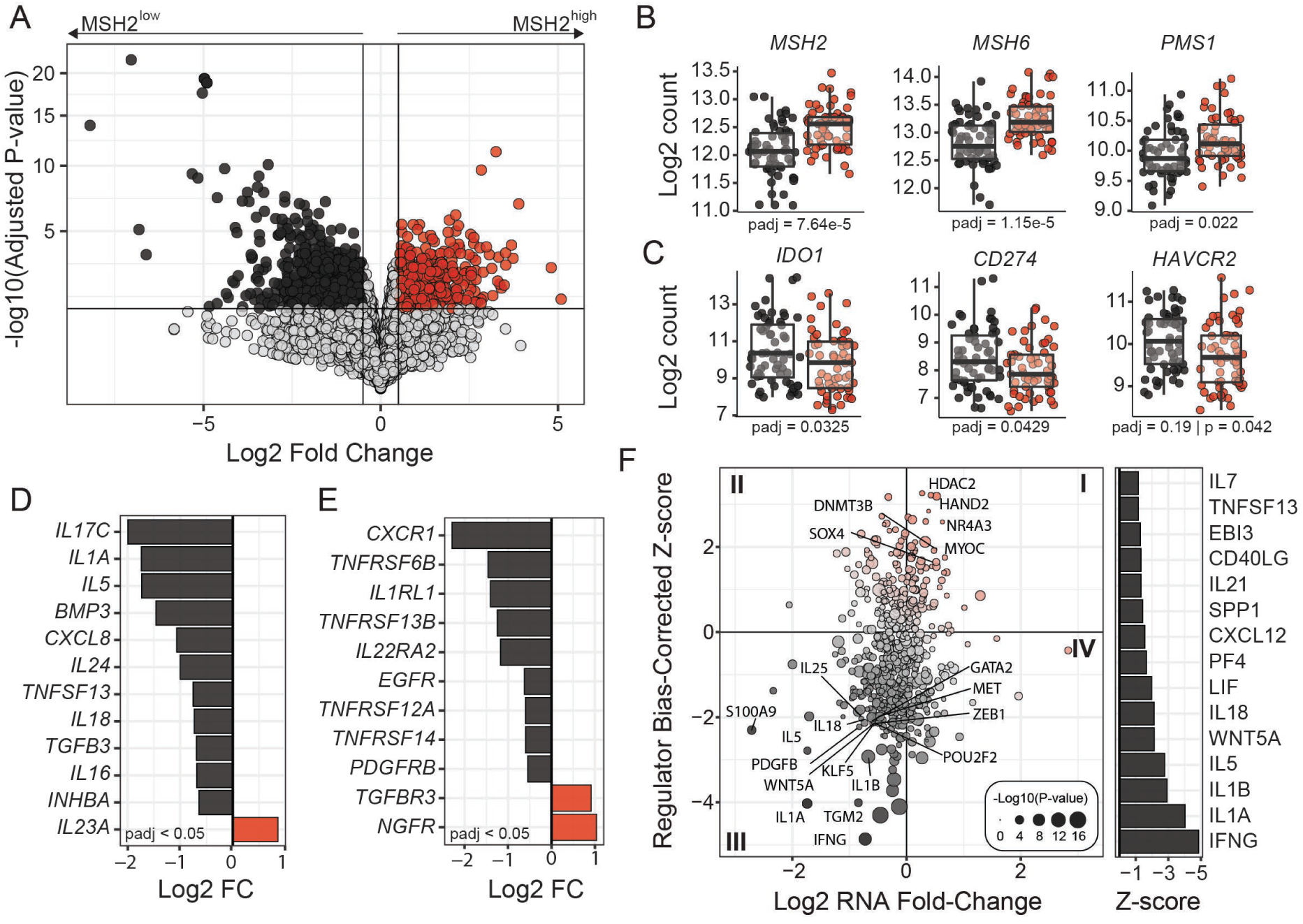
Differential gene expression results based on MSH2 protein in the TCGA BLBC patient specimens. **A.** Differential gene expression using HTseq count data comparing MSH2-high (n=60) to MSH2-low (n=59) found 650 genes significantly increased and 1,444 significantly decreased. Significance was defined as an adjusted P-value < 0.05 and log2 fold-change ≥ |0.5|. **B.** Regularized log2 transformed count expression of DNA mismatch repair pathway constituents that demonstrated significantly differential between MSH2-high versus low. **C.** Regularized log2 transformed count expression of immune checkpoint targets. **D.** Log2-fold change of significant chemokines and cytokines. **E.** Log2-fold change of significant chemokine and cytokine receptors. **F.** IPA upstream regulators results based on differential gene expression. Quadrant I correspond to regulators with increased expression and predicted increase activity in MSH2-high samples, while Quadrant III corresponds to increased expression and predicted activity in MSH2-low samples. Bar graph summarizes the significant predicted activation of cytokines.

**Figure S11.**
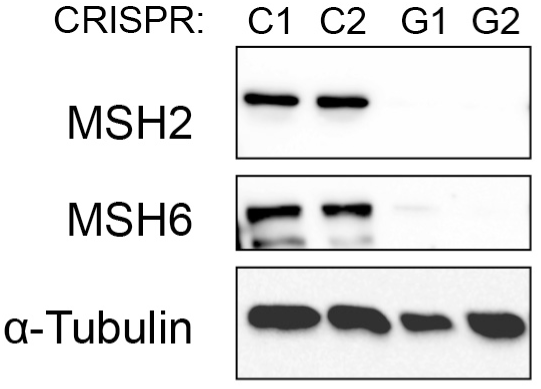
Generation of MSH2 KO MDA-MB-231 BLBC cells. Lentiviral particles encoding Cas9 and guide RNA for MSH2 were used to infect MDA-MB-231 cells, following 5-day selection process with puromycin to induce MSH2 KO. The MSH2 KO cells were passaged to P30 and KO efficiency was confirmed using immunoblotting, with concomitant loss of MSH6.

